# Evolution of division of labor on two public goods with continuous investments

**DOI:** 10.1101/2025.03.27.645589

**Authors:** Aidan Fielding, Erol Akçay, Joshua Plotkin

## Abstract

Species throughout the tree of life have evolved to produce multiple public goods, and they often exhibit division of labor, meaning that subpopulations have different allocations of effort across the goods. Despite a robust theoretical literature on mechanisms that lead to division of labor in specific domains, a general evolutionary game-theoretical analysis remains incomplete. Here we model labor as a continuous quantity and the resulting public goods as functions of labor contributed by all individuals using adaptive dynamics. We derive general conditions for the evolution of division of labor and categorize what forms of division may arise. We show that the adaptive dynamics of division of labor has three potential outcomes that differ in expected payoff and variance in payoff. Our findings indicate that the evolution of division of labor can have a diverse range of possible outcomes, with important consequences for fitness, variance in payoff, and the distribution of effort across the population.

## Introduction

Stable patterns of cooperation are an important part of the ecology of diverse species, from microbes to humans (Hamilton, 1964; Maynard Smith and Price, 1973; Axelrod and Hamilton, 1981; Frank, 1995; Apicella and Silk, 2019). For example, many bacteria rely on production of public goods like extracellular enzymes (Smith and Schuster, 2019), biofilm substrates (Drescher et al., 2014), or iron-scavenging siderophores (Cordero et al., 2012; Kramer et al., 2019) to effectively grow colonies. These public goods are typically costly to those who produce them, but once they are produced all individuals can take advantage, regardless of how much each individual contributed (Nowak, 2006; Archetti and Scheuring, 2012). This creates a well-known social dilemma where public goods might end up being under-supplied relative to the level that would be socially optimal, because of the incentive to free-ride (Rankin et al., 2007; Hauert et al., 2007; Frank, 2010; Akçay and Van Cleve, 2012).

Such public goods dilemmas abound in microbial ecology: an illustrative example is siderophore production in some bacterial populations (Smith and Schuster, 2019; Cordero et al., 2012; Kramer et al., 2019). Each bacterium ‘invests’ part of its metabolic activity into producing siderophores and secreting them into the extracellular environment, paying a fitness cost that is a function of this metabolic activity. The siderophores diffuse into the environment and facilitate the uptake of iron by both the focal individual and its neighbors. Another example is production and secretion of of invertase, an enzyme that hydrolyzes sucrose, by yeast growing on sucrose, where most hydrolysis products diffuse away from the focal cell and therefore benefit other cells in the culture (Gore et al., 2009).

In many such cases, the production of the benefits is non-linear, such that the public good can be produced when only a subset of individuals invest into it. These situations give rise to a class of strategic interactions called snowdrift games or volunteer’s dilemmas (Maynard Smith and Price, 1973; Diekmann, 1985; Hauert and Doebeli, 2004). In such social dilemmas, an evolutionarily stable outcome often involves two types investing unequal amounts, which can be considered an example of division of labor.

Investments in a public good can be modeled as a discrete strategy (i.e., cooperate or defect; Archetti and Scheuring, 2012), as mixed strategies depicting the probability of taking a discrete strategy (Motro, 1991; Zheng et al., 2007; Souza et al., 2009), or as a continuous quantity depicting the *level* of investment into the public good (Dieckmann and Law, 1996; Doebeli et al., 2004; Henriques et al., 2021). In each of these contexts, conditions have been derived that determine whether a population will evolve to a dimorphic state, with some cooperators (or, high-investors) and some defectors (low-investors). When investment is described by a continuous quantity, the population may transition from a monomorphic state, with equal investment levels for all, to a dimorphic state containing sub-populations of high- and a low-investing types (Doebeli et al., 2004). This process of evolutionary branching into two types is a one-dimensional example of emergent division of labor, in which novel investment types evolve and coexist.

In real populations, there is often more than one public good being produced and consumed. For example, some bacteria rely on production of multiple types of metabolites to facilitate colony growth: *P. aeruginosa* colonies require a siderophore and an elastase, both of which are public goods, to successfully expand (Ross-Gillespie et al., 2014; Özkaya et al., 2017). Additionally, in cancer cells, production of acid through glycolysis and facilitation of oxygen absorption through vascularization are shared goods that enable metastasis (Kaznatcheev et al., 2017). In such cases, individuals who are ‘cooperators’ for one good (meaning, they pay the cost of production) may be ‘defectors’ for the other good (meaning, they do not contribute to production) (Oliviera et al., 2014; Rossetti et al., 2010; West and Cooper, 2016; Cooper and West, 2018).

Likewise, human hunter-gatherer groups sustain themselves by cooperatively foraging for both meat and wild plants (Apicella and Silk, 2019; Stibbard-Hawkes et al., 2022). Group members allocate their effort across both tasks, and nutritional benefits are often shared with non-foraging members. The payoff to each individual from each public good is equal to the collective benefit they receive from everyone’s foraging effort minus the individual cost they pay for their own effort.Notably, hunter-gatherer groups often divide labor such that some individuals devote more effort to hunting for meat while others invest more in gathering plants (Bird, 1999; Hooper et al., 2015; Anderson et al., 2023). While this pattern is often attributed to behavioral differences between sexes in human populations, it remains important to ask whether such patterns may arise in the absence of any *a priori* differences among individuals. More generally, we seek to understand whether the emergence of division of labor will always lead to outcomes like this, involving specialization on one good or another, or whether other forms of labor structures are possible.

With two public goods, three distinct forms of division of labor are known to exist (Henriques et al., 2021). In the first form, some individuals invest more in both goods while others invest less in both goods (“Exploitation,” which we abbreviate as “ex”); in the second, some individuals invest more in good one and less in good two, while others invest more in good two and less in good one (“Specialization,” abbreviated as “sp”) (Doebeli et al., 2004; Henriques et al., 2021). Previous work by Henriques et al. (2021) in two-player games showed that depending on the initial genetic variation, populations might evolve to either outcome. Here, we use adaptive dynamics to study the emergence and consequences of these distinct forms of division of labor, and a novel third form, in multiplayer public good games with two goods.

Specifically, we consider the evolution of continuous investment levels in two public goods, where an individual’s payoff is a function of the collective benefit derived from all group members’ investments as well as the personal cost of an individual’s own investments. If the payoff function allows for evolution of division of labor on both public goods, we show that the population might evolve to either the exploitation or the specialization outcome, depending on the initial distribution of traits. We then characterize the evolutionary dynamics of these two outcomes and compare the payoff consequences for individuals in populations that took either path. Notably, we find that variance in payoff from the public goods is higher in the exploitation than in the specialization forms of labor division; but mean fitness is also marginally higher in the exploitation outcome. Differing payoff variance can reshape dynamics compared to classical models that depend only on fitness differences (Gillespie, 1974, 1975; Constable et al., 2016; Wang et al., 2023). We also show that both exploitation and specialization populations can evolve yet more complex forms of labor division, and we analyze a third intermediate form in which investments in one of the goods are constant across types. These results highlight the wide range of evolutionary outcomes that can arise in populations producing and consuming multiple public goods.

## Methods

To build intuition, we use siderophore production in bacterial populations as an example of a single public good. Individuals in a colony receive a public benefit that is a function of their own investment into production and their neighbors’ investments. They pay a cost that is a function only of their own investment. A cluster of *n* cells can be represented by a group of *n* players, with player *i* ∈ (1, …, *n*) investing amount *z*_*i*_ ∈ ℝ^+^ in the public good. The benefit *i* receives, *B*(*z*_1_, *z*_2_, …, *z*_*n*_), is a function of all *n* investments. The cost that *i* pays, *C*(*z*_*i*_), is a function of *i*’s investment. The benefit and cost functions *B* and *C* are assumed to be non-negative, continuous, and increasing with the level of investment. The payoff that *i* receives from the public goods game is written π(*z*_*i*_, ⋅) = *B*(*z*_*i*_, ⋅) − *C*(*z*_*i*_), where ⋅ denotes the other *n* − 1 investments (not *i*’s).

### Adaptive dynamics of investments in one public good

#### One type

We assume that the population is large, and that individuals play *n*-person public goods games with *n*−1 randomly drawn others from the population. We consider a rare mutant with investment *z*_mut_, in an otherwise monomorphic population with resident investment level *z* (Dieckmann and Law, 1996; Metz et al., 1996). The mutant’s fitness is equal to the expected payoff it receives from the public goods game:

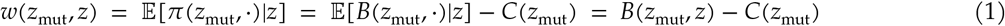

Where we have denoted the expected value of the public benefit for the mutant as 𝔼 [*B*(*z*_mut_, ⋅)|*z*] = *B*(*z*_mut_, *z*), that is, the benefit received by the mutant when all *n* − 1 other players invest *z*. Below, we make the abbreviation *B*(*z*_mut_, *z*) ∶= *B*.

Under the framework of adaptive dynamics, the resident investment evolves in the direction of the selection gradient:

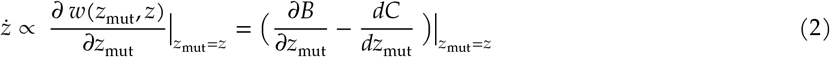

A critical point *z*_∗_ of the dynamics occurs when the selection gradient is zero:

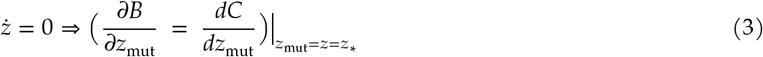

Evolution of the resident investment *z* near a critical point *z*_∗_ can be classified into three different regimes (Geritz et al., 1998; Doebeli et al., 2004) (also see Figure 2). The simplest possibility is that *z*_∗_ is convergently and evolutionarily stable, so that a monomorphic population with resident investment *z* in a neighborhood of *z*_∗_ will evolve to and equilibrate at *z*_∗_. Alternatively, *z*_∗_ can be convergently and evolutionarily unstable, so a resident investment *z* in a neighborhood of *z*_∗_ will evolve away from *z*_∗_. Lastly, there could be evolutionary branching that produces a stable dimorphism near *z*_∗_, which is only possible if the mutant fitness function *w*(*z*_mut_, *z*) meets the following conditions (Geritz et al., 1998; Leimar, 2009):

**Table 1.** Evolutionary branching conditions for various model families

|  | Payoff function | Branching conditions |
| --- | --- | --- |
| General | $\pi_i = B(f(z_i, \cdot)) - C(z_i)$ | $\left. \frac{d^2 B}{df^2} \right\vert _* < 0$ and $\left. \frac{d^2 C}{dz_{\text{mut}}^2} \right\vert _* < 0$ |
| Summed $z$ | $\pi_i = B\left(\sum_{j=1}^n z_j\right) - C(z_i)$ | $\left( n \frac{\partial^2 B}{\partial z_{\text{mut}}^2} < \frac{\partial^2 C}{\partial z_{\text{mut}}^2} < \frac{\partial^2 B}{\partial z_{\text{mut}}^2} < 0 \right) \Big\vert _*$ |
| Multiplicative $z$ | $\pi_i = B\left(\prod_{j=1}^n z_j\right) - C(z_i)$<br>$(f(z_i, \cdot) = \prod_{j=1}^n z_j)$ | $\left( n z_*^{2(n-1)} \frac{d^2 B}{df^2} + (n-1) z_*^{n-2} \frac{dB}{df} < \frac{d^2 C}{df^2} < z_*^{2(n-1)} \frac{d^2 B}{df^2} < 0 \right) \Big\vert _*$ |
| Model 1 | $\pi_i = g(\sum z_j)^{a_1} - z_i^{a_2}$ | $a_1 - 1 < a_2 - 1 < (a_1 - 1)/n$ |
| Model 2 | $\pi_i = -a_1(\sum z_j)^\alpha + a_2(\sum z_j) - (-a_3 z_i^\alpha + a_4 z_i)$ | $-a_1 n^{\alpha-1} < -a_3 < -a_1 n^{\alpha-2}$ |

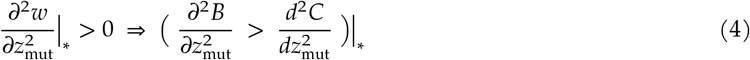

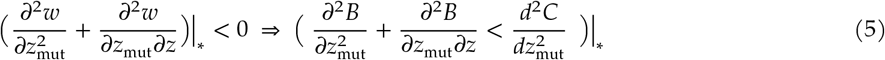

Where “*” indicates *z*_mut_ = *z* = *z*_∗_, that is, the mutant trait is set equal to the resident trait *z* which is at the monomorphic critical point *z*_∗_.

Evolutionary branching producing dimorphic effort levels can only occur if the benefit and cost functions satisfy conditions (4), which confers evolutionary instability, and (5), which confers convergence stability. Note that (4) and (5) cannot both be true if 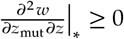. Note also that if inequality (4) is false while (5) is true, the monomorphic critical point is convergently and evolutionarily stable, with everyone investing the same positive amount in the public good.

#### Two types

After branching, two resident types co-exist in the population. We label them type *u* and type *v*, with investment levels *z*_*u*_ and *z*_*v*_. The frequency of type *u* in the population is *p* ∈ [0, 1], and the frequency of type *v* is 1 − *p*. Following the same derivation as for one good, fitness for a mutant with trait 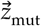 is equal to the expected payoff when its *n* − 1 opponents in the public goods game are drawn from the population with resident types *u* and *v* with probabilities *p* and 1 − *p*, respectively. That is

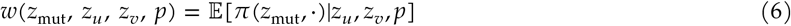

One of the assumptions of adaptive dynamics is a separation of timescales such that resident types will reach their frequency equilibrium *p*_∗_ before any new mutant is introduced. At *p*_∗_, the fitnesses of types *u* and *v* are equal:

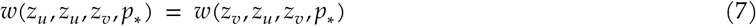

The two-type selection gradient is computed by taking the partial derivative of fitness with respect to the mutant trait and evaluating at *z*_mut_ = *z*_*u*_ (for type *u*) and *z*_mut_ = *z*_*v*_ (for type *v*) when the resident traits *z*_*u*_ (*z*_*v*_) are at equilibrium frequencies *p*_∗_ (1 − *p*_∗_):

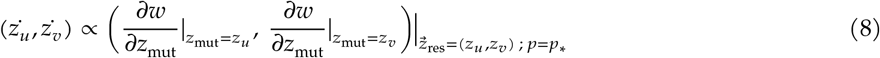

Adaptive dynamical trajectories depend only on differences in fitness (expected payoff) and are not affected by differences in payoff variance between resident types. However, it has been shown that differences in payoff variance can alter evolutionary dynamics from those that depend on fitness alone (Gillespie, 1974, 1975; Constable et al., 2016; Wang et al., 2023). For an arbitrary individual *q* with investment *z*_*q*_ in a population with resident types *u* and *v*, we can compute *q*’s variance in payoff as follows:

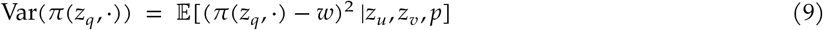

### Adaptive dynamics with two public goods

We consider the simplest case where the benefits from the two public goods, and the costs of investing into them, are additive (Figure 1). Each individual now has two investment traits. For individual *i* ∈ (1, …, *n*), their investments are written 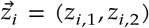, where the first subscript indicates the player ID and the second subscript indicates the public good (either 1 or 2). In this setting, there is the possibility for division of labor in the sense that different portions of the population may allocate their effort differently across the two goods.

**Figure 1:**
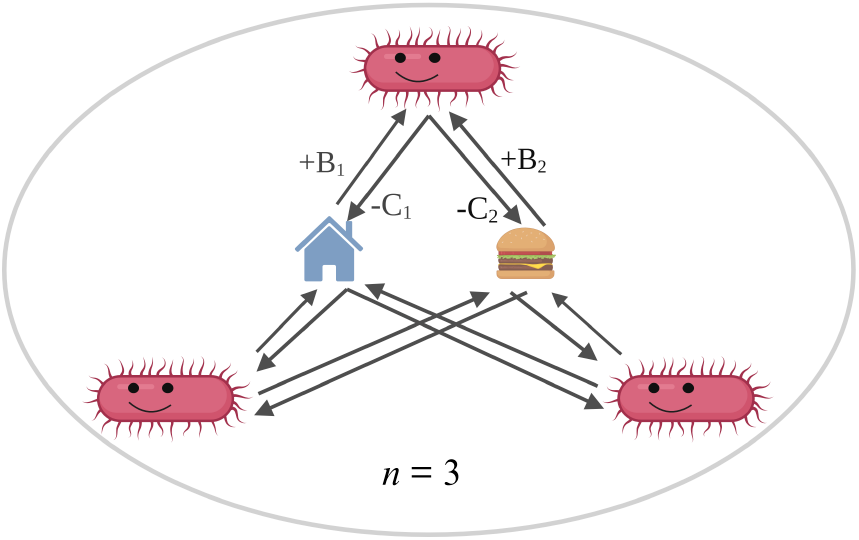
Illustration of a public goods game with 2 goods and *n* = 3 players. Here, 3 bacteria are investing in 2 public goods. Good one is biofilm production (‘shelter,’ represented by a house), with public benefit *B*1 and private cost *C*1. Good two is the facilitation of iron uptake via siderophores (‘food,’ represented by a cheeseburger), with benefit *B*2 and cost *C*2.

**Figure 2:**
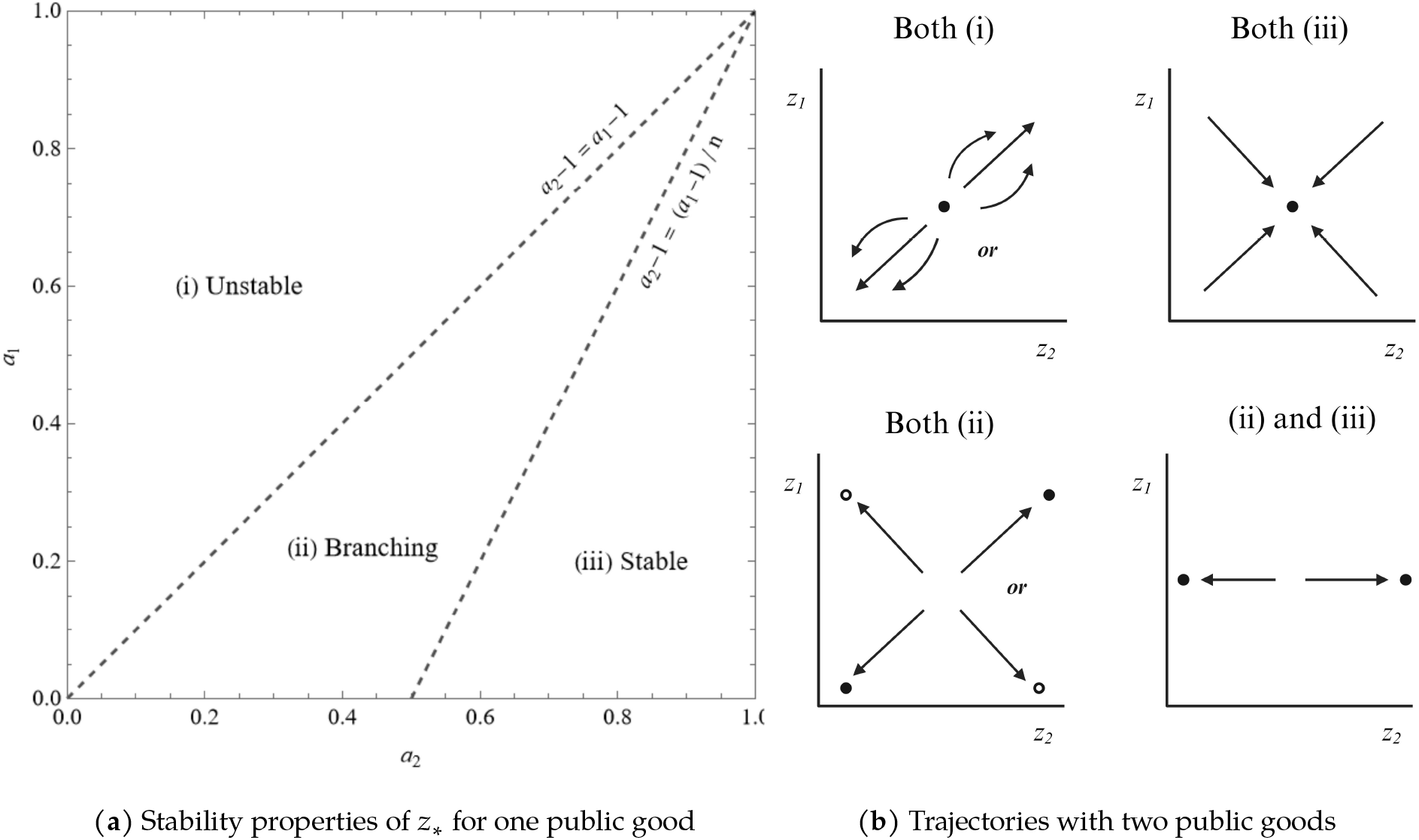
Adaptive dynamics of investments for model 1, with *n* = 2. (a) Evolutionary stability properties of the monomorphic critical point *z*_∗_ with a single public good (see model 1, table 1) across parameter space. The axes are *a*_1_ and *a*_2_, which are parameters that control the concavity of the public benefit and private cost functions. (*i*) In the unstable region, *z*_∗_ is convergently unstable and evolutionarily unstable, so a population starting above (below) *z*_∗_ will evolve upward (downward). (*ii*) In the branching region, *z*_∗_ is convergently stable but evolutionarily unstable, so a monomorphic population will evolve to *z*_∗_ and then branch into two types. (*iii*) In the stable region, *z*_∗_ is convergent and evolutionarily stable, so a monomorphic population will evolve to *z*_∗_ and stay there without branching. (b) Subplots indicate the adaptive dynamics of investments into two public goods when their respective parameters are in the given regions from the left plot. In each subplot, the vertical (horizontal) axis represents investment level in good one (good two), which we denote *z*_1_ (*z*_2_). In the top row, the population is monomorphic (one type). If *a*_1_, *a*_2_ (*a*_3_, *a*_4_) are in the unstable region (top-left), the resident investment level will evolve away from the monomorphic critical point, while if they are in the stable region (top-right), the monomorphic critical point is the ESS. In the bottom row, the population is dimorphic (two types) after evolutionary branching. If *a*_1_, *a*_2_ (*a*_3_, *a*_4_) are both in the branching region (bottom left), dynamics vary with initial conditions: the dimorphic population that emerges due to branching evolves to one of two critical points, the exploitation outcome (filled circles) or the specialization outcome (empty circles). If *a*_1_, *a*_2_ are in the stable region but *a*_3_, *a*_4_ are in the branching region (bottom right), the dimorphic population divides its labor on good two, but maintains a constant investment level on good one.

Let *B*_*k*_ and *C*_*k*_ denote the benefit and cost functions for good *k*, and let {*z*_−*i,k*_} denote the *n* − 1 nonfocal investments in good *k*. Payoff for *i* in this two public good setting can be written as follows:

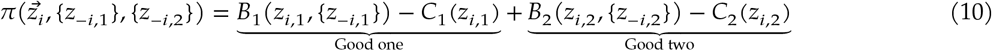

Note that in this simple setting, we assume no interactions between the public goods, so the public benefit function for each good does not take investments into the other good as inputs, and the payoffs from both goods are summed to get total payoff.

If (4) and (5) are true for either good one (*B*_1_, *C*_1_) or good two (*B*_2_, *C*_2_), then evolutionary branching into two types will occur. If (4) and (5) are true for both goods, evolution of the two types can result in two characteristically different divisions of labor: exploitation (ex), in which some individuals invest more in both goods while others invest less in both; and specialization (sp), in which some individuals invest more in good one and less in good two, while others invest more in good two and less in good one (Figure 2).

As before with one public good, we label the two types that arise after branching *u* and *v*, with investment traits 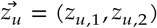 and 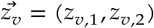 and frequencies *p*, (1 − *p*). The fitness (50) of a subsequent mutant is is given by its expected payoff from the public goods, with the trait values replaced by the appropriate trait vectors. The frequency equilibrium *p*_∗_ (7), the selection gradients of the types’ investments (8), and the variances in payoff of types (9) are defined similarly.

## Results

### Evolutionary branching in a single good

We start by analyzing the adaptive dynamics of investment levels in a single public good. First, we define a set of intuitive assumptions about the public benefit function *B* and the private cost function *C* and show that under these assumptions, *B* and *C* must both have diminishing returns to their investment inputs in order for evolutionary branching to occur.

Consider the general payoff function π(*z*_*i*_, ⋅) = *B*(*z*_*i*_, ⋅) − *C*(*z*_*i*_). We first assume that *B* and *C* are non-negative, continuous, and increase with investments, 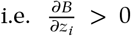 and 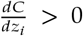 for all *i* ∈ (1, …, *n*). Second, we assume that *B* is a composite function of all investments. Payoffs can then be denoted π(*z*_*i*_, ⋅) = *B*(*f* (*z*_*i*_, ⋅)) − *C*(*z*_*i*_), where 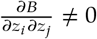 for all *j* ≠ *i* ∈ (1, …, *n*). The key property is that the cross-derivative of focal and non-focal investments is nonzero, which is required for evolutionary branching (see condition (5)). Conceptually, the composite nature of *B* represents a group-level effect of public good production. Note also that 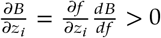 implies that 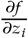 and 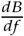 must have the same sign.

Next, we assume that individuals’ contributions to the public good are *exchangeable*, i.e., one individual reducing their investment while another increases by the same amount does not change the overall benefit produced, and the same benefit and cost functions apply for all individuals. This represents a null hypothesis that individuals do not vary in their competence at contributing to the public good. Technically, this property only needs to hold when investments are at the monomorphic critical point *z*_∗_, because that is the only point at which evolutionary branching can occur. Referring to the inner function *f* of the composite benefit function *B*(*f* (*z*_*i*_, ⋅)), exchangeability implies that 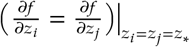 for all *i, j* ∈ (1, …, *n*).

Third, we assume that the cross-derivative 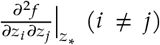 has the same sign as 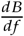 and 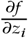 In words, the second-order effect of each individual’s investment on others’ investments has the same sign as each individual’s first-order effect on public benefit. Lastly, we assume that 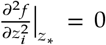, meaning that any nonlinear effect of focal investment on public benefit is due to the outer function *B*.

We can show (Appendix A.2) that if *B* and *C* meet the assumptions defined above, the following conditions are necessary for a monomorphic critical point *z*_∗_ to be an evolutionary branching point:

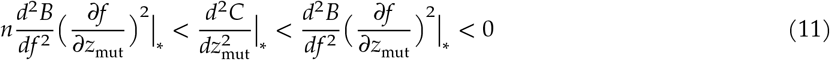

One implication of this result is that the evolution of division of labor by small mutations in a well-mixed population requires that both public benefits and private costs have diminishing returns with respect to investments in the goods. Another implication is that if 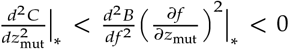, then increasing the number *n* of players in the public goods game will ensure that *z*_∗_ is an evolutionary branching point (above some threshold value).

Table 1 lists evolutionary branching conditions for several families of *B* and *C* functions at varying levels of generality, all of which meet the assumptions above. Payoff functions are written for focal player *i*, and the derivation proceeds from (4) and (5). Numerical models 1 and 2 are subfamilies of the “Summed *z*” family defined in row 2. Note that for the Summed *z* family, 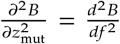. The stability properties of *z* for model 1 (table 1, row 3) across parameter space are depicted in Figure 2a; evolutionary branching conditions are satisfied in region (*ii*). Model 2 (table 1, row 4) is equivalent to a payoff function studied by Henriques et al. (2021) in a similar context.

The functional conditions for the Summed *z* family and the parametric conditions for models 1 and 2 all express that 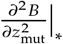 and 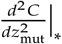 must be negative and fall within a relative range that grows with *n*. If either of the first two inequalities in the Summed *z* conditions are violated, polymorphism never emerges. Figure 2b (top row) depicts the different regimes of evolutionary dynamics that can occur depending on which inequality is violated (Doebeli et al., 2004).

After evolutionary branching, two resident types *u* and *v* exist in the population with frequencies *p* and 1 − *p*. When public benefits are a function of the summed investments (table 1, row 2), we can derive unique adaptive dynamical trajectories of both types’ investments if and only if benefit function *B* exhibits diminishing returns to its inputs. This is because if a critical frequency 0 < *p*_∗_ < 1 solving (7) exists for *u* and *v*, it will be unique and dynamically stable if and only if 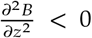 for *z* ≥ 0 (we show this in Appendix B.1 using Result 3 from Peña et al., 2014). The assumption of diminishing returns for *B* on its entire domain is stronger than only assuming this concavity at the critical point, but both represent a saturation constraint on public goods production. Note that this condition is satisfied in model 1 whenever evolutionary branching conditions are satisfied.

### Evolutionary branching in two goods

If the payoffs to individuals from two public goods are additive, as in (10), the conditions on *B*_*k*_ and *C*_*k*_ for evolutionary branching of investments in good *k* ∈ {1, 2} are equivalent to the conditions obtained from analyzing good *k* in isolation. Thus, in the two good setting, if at least one public good meets branching conditions, a dimorphic population will evolve. If both goods meet branching conditions, either the exploitation or the specialization division of labor will evolve. These dimorphic outcomes are illustrated in Figure 2b (bottom row). Which outcome evolves depends on the stability properties of both goods analyzed in isolation, which are shown in Figure 2a for model 1, as a function of *a*_1_ and *a*_2_.

The exploitation and specialization divisions of labor (subplot (b), bottom left) occur when the parameters for both goods are in subplot (a) region (*ii*), implying evolutionary branching for both goods in isolation. An intermediate form of division of labor (subplot (b), bottom right) occurs when the monomorphic critical point for one public good is a convergently stable ESS, while that of the other good is an evolutionary branching point. With model 1, this occurs if the parameters for one good are in subplot (a) region (*iii*) and parameters for the other good are in region (*ii*). Like the exploitation outcome, this third configuration is qualitatively similar to dimorphic populations in the one-good setting, in that there is a cooperator and a defector type.

### Evolution of division of labor

Next, we numerically integrate the adaptive dynamics of investments in two public goods after evolutionary branching. To do this, we assume that the payoffs from the goods are additive, as in (10), and that the payoffs from either good in isolation have the functional form of model 1 (table 1, row 3):

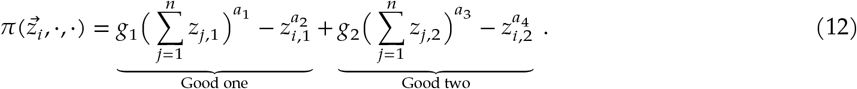

The parameters *a*_1_ and *a*_3_ govern the concavity of the benefit functions for goods 1 and 2, respectively, while *a*_2_ and *a*_4_ govern the concavity of the cost functions for goods 1 and 2, respectively. The quantities *g*_1_ and *g*_2_ are scaling parameters that do not affect evolutionary stability.

The following four sections present the comparative statics of the exploitation and specialization divisions of labor for model 1 with two public goods. Specifically, we present the investment levels, fitness, payoff variance of both types, and mean payoff variance, evaluated at the critical points of adaptive dynamics with two types. In other words, we compute when the resident population (*z*_*u*,1_, *z*_*u*,2_, *z*_*v*,1_, *z*_*v*,2_, *p*_∗_) has selection gradient equal to zero for all four investment traits. This is the final point in the expected trajectory for two types, after which the population will either remain in the same state or possibly undergo further branching (Leimar, 2009). Evaluating our quantities of interest at the respective dimorphic critical points for the exploitation and specialization outcomes is thus an intuitive way to compare them. We present results below with *n* = 2 and identical goods (*a*_1_ = *a*_3_, *a*_2_ = *a*_4_, *g*_1_ = *g*_2_ = 2) which therefore have the same monomorphic critical point *z*_∗,1_ = *z*_∗,2_ = *z*_∗_ when analyzed separately. We have verified that qualitative relationships hold for *n* = 3, other values of *g*_1_ and *g*_2_, and small values of |*a*_1_ − *a*_3_| and |*a*_2_ − *a*_4_|.

### Investment levels and type frequencies for exploitation and specialization divisions of labor

The distributions of investments in the exploitation and specialization division of labor are shaped by the relative concavity of public benefits *B* and private costs *C*. For values of *a*_1_ and *a*_2_ that meet evolutionary branching conditions, the adaptive dynamics of 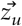 and 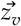 either have four or eight dimorphic critical points, each of which can be classified as exploitation or specialization. Investment levels and type frequencies at a representative subset of these points are plotted in Figure 3.

**Figure 3:**
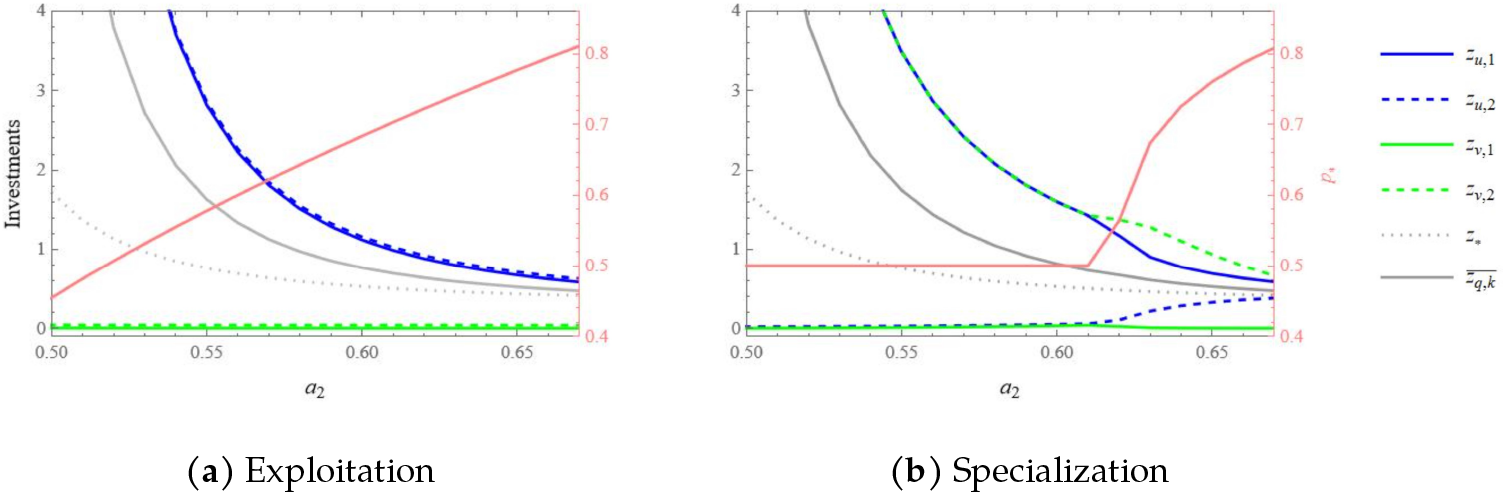
Critical point investments and frequencies for both outcomes. In this plot, *a*_1_ = *a*_3_ = 0.4, *a*_2_ = *a*_4_, *n* = 2, and *g* = 2. In both cases, as *a*_2_ increases, the upper investment levels, the mean investment level 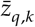, and the monomorphic critical point *z*_∗_ all decrease and converge on each other. With exploitation, this is accompanied by a compensatory increase in the frequency *p*_∗_ of the high-investing type (*u*). With specialization, there is a bifurcation when *a*_2_ is sufficiently high. Below the threshold value of *a*_2_, investments across goods are symmetric and frequencies are equal (*p*_∗_ = 0.5). Above the threshold, the upper and lower investments of one type (*u*, above) evolve towards each other and approach *z*_∗_. The frequency of this type increases above 0.5 as *a*_2_ increases further, due to the large marginal public benefits produced by this type increasing its lower investment. The upper and lower investments of the other type (*v*, above) evolve away from each other after the bifurcation in *a*_2_ but then approach each other as *a*_2_ increases. Note that for all dimorphic critical points except specialization points that are asymmetric across goods, the mean investment levels in both goods are the same; in the asymmetric-specialization region of subplot *b*, the grey line depicts the mean investment into good one.

As *a*_2_ increases, the marginal private cost of investment diminishes slower (becomes closer to linear) relative to the public benefit. This makes the evolutionary branching point *z*_∗_ decrease. Simultaneously, the dimorphic critical points come closer to the branching point, so the difference between the types in the division of labor outcomes becomes smaller. Intuitively, this is because branching and diversification is favored by strongly diminishing marginal costs. Summarizing these effects, the mean investment level *p*_∗_*z*_*u,k*_ + (1 − *p*_∗_)*z*_*v,k*_ into good *k* ∈ {1, 2} in the dimorphic population (Figure 3, solid grey line) decreases and approaches *z*_∗_ (Figure 3, dotted grey line) as *a*_2_ increases.

All exploitation critical points (Figure 3a) and a majority of specialization critical points (Figure 3b, left) have equal investment levels for public goods 1 and 2 (though levels differ across these two outcomes). In an exploitation outcome, the “cooperator” type invests an amount greater than *z*_∗_ in both goods while the “defector” type invests an amount less than *z*_∗_ in both. If *u* is the cooperator type, we have *z*_*v*,1_ = *z*_*v*,2_ < *z*_∗_ < *z*_*u*,1_ = *z*_*u*,2_. In a specialization outcome, each specialist type invests greater than *z*_∗_ in one good and less than *z*_∗_ in the other. If *u* specializes on good one, for example, we have *z*_*v*,1_ = *z*_*u*,2_ < *z*_∗_ < *z*_*u*,1_ = *z*_*v*,2_. Multiplying by two possible type-orderings, there are four critical points for each parametrization when this symmetry across goods holds.

When *a*_2_ sufficiently exceeds *a*_1_, the specialization outcome has four additional dimorphic critical points that have asymmetric investments across goods 1 and 2. One of these points is plotted in Figure 3b: at this point, (*z*_*v*,1_ < *z*_*u*,2_) < *z*_∗_ < (*z*_*u*,1_ < *z*_*v*,2_), so type *u* still specializes on good one but more of good two is produced overall (the three other asymmetric points are listed in Appendix C.4).

Overall, the difference between a multistable pair of exploitation and specialization divisions of labor depends on how fast the marginal private cost of investment diminishes relative to the marginal public benefit.

### Population fitness for exploitation and specialization outcomes

The exploitation outcome – in which one type provides higher investment in both goods – might seem to be an “unfair” division of labor, compared to the specialization outcome. At the frequency equilibrium *p*_∗_, the expected fitness of coexisting resident types must be equal by definition (equation (7)), but this expected fitness might differ between the two outcomes. Therefore, we next evaluate the fitness consequences of the two forms of labor division.

We compare resident fitness in the exploitation outcome (*w*_*ex*_) to that in the specialization outcome (*w*_*sp*_) (Figure 4). For every parametrization of model 1 that allows evolutionary branching, we find that the exploitation outcome produces greater resident fitness, that is *w*_*ex*_ > *w*_*sp*_. We also numerically compute the value of *a*_2_ at which △*w* = *w*_*ex*_ − *w*_*sp*_ reaches a local maximum. That value coincides with the value of *a*_2_ at which specialized dimorphisms with asymmetric investments across goods arise (Figure 3b). This gives us an intuition for why these asymmetric dimorphisms emerge: below the threshold value of *a*_2_, specialized types have symmetric investments across goods, and their frequencies are therefore constrained to be (0.5, 0.5). But as *a*_2_ increases (i.e., the costs of investment becomes closer to linear), upper investment levels decrease, thus obtaining a higher marginal benefit. In the exploitation outcome, this is accompanied by a compensatory increase in the frequency of the cooperating *u* type, but in the specialization outcome there are no frequency changes before the bifurcation. This results in a fitness deficit that accumulates until a threshold value of *a*_2_, beyond which the specialists can break symmetry and improve their fitness relative to the corresponding exploitation population.

**Figure 4:**
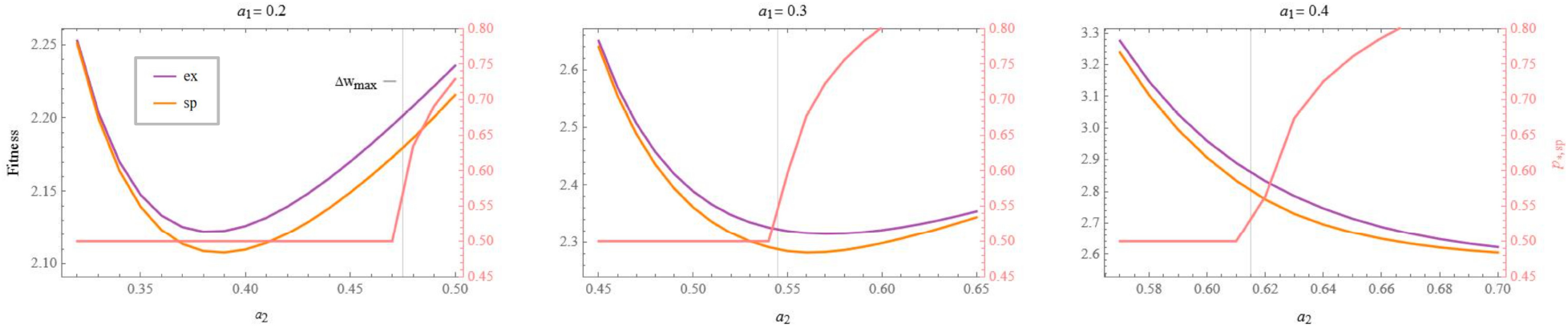
Fitness for both outcomes and *p*_∗_ for specialization. The purple and yellow lines show the fitness of an individual in the exploitation and specialization outcomes, respectively, depending on *a*_2_. The value of *a*_2_ that maximizes the difference △*w* = *w*_ex_ − *w*_sp_ is indicated by a grey vertical line. The pink line in each plot shows the frequency equilibrium *p*_∗,sp_ for the specialization outcome. The value of *a*_2_ at which *p*_∗,sp_ increases above 0.5 coincides with that which maximizes △*w*.

We have shown that fitness is higher for exploitation than for specialization divisions of labor, due to the essential difference between their distributions of investments across goods. This result favors exploitation-type populations over their specialized counterparts, but in the next section we show that patterns of payoff variance across the two outcomes favor specialist populations.

### Variance in payoff across types and across outcomes

Another divergent consequence of the two kinds of division of labor is that while individuals in a specialization outcome can always rely on some of each public good being produced, this is not true for the exploitation outcome. This suggests individuals in the exploitation outcome might experience more variance in their payoff. Demographic stochasticity due to such variance in payoff can influence the dynamics of competing types (Gillespie, 1974; Constable et al., 2016; Kennedy et al., 2018; Wang et al., 2023). Therefore, we next consider the patterns of payoff variance in the two outcomes.

We calculate the payoff variance of types *u* and *v* 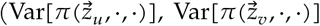; see (9) and Appendix C.1) and the mean variance in the population 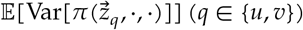 for both the exploitation and specialization outcomes, for all parametrizations of model 1 that allow evolutionary branching (Figure 2a, region (*ii*)). Three qualitative patterns are consistent across all parametrizations. First, with *one* public good and two types (cooperators and defectors), the defecting type has much higher variance. Second, with *two* public goods, the mean variance in the exploitation outcome is much higher than that in the specialization outcome (Figure 5 row 2). Third, the ranking of payoff variance among the four types changes as *a*_1_ increases (Figure 5 row 1). For low *a*_1_, the variance of the cooperating type is below that of either specialized type, which in turn is well below that of the defecting type. With higher *a*_1_, the variance of the cooperating type exceeds that of specialists, and approaches that of the defectors.

**Figure 5:**
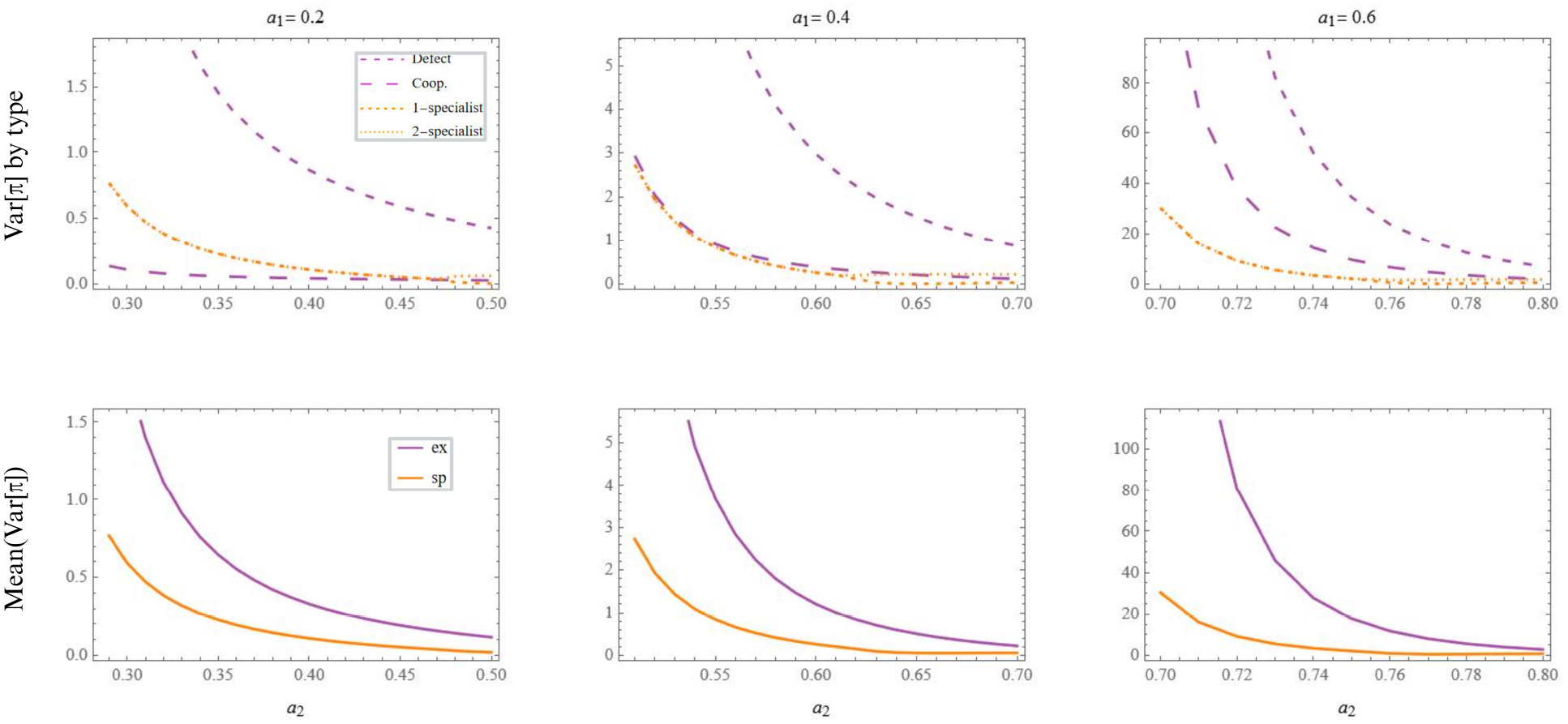
Variance in payoff across types and outcomes. Row 1 shows the variance in payoff for each type (9) for parameters *a*_1_ and *a*_2_. Row 2 shows the mean payoff variance for either outcome. Other parameters for the model specifications in this plot are *n* = 2 and *g* = 2.

In general, the defecting type has the highest payoff variance, for either one or two goods. This is because of the diminishing returns to investments in the public benefit (which are required for the types to coexist). Consider the case when *n* = 2. For both goods, a defector’s opponent will either defect or cooperate. The difference in pay-offs between these two scenarios corresponds to the first ‘margin’ of the benefit function, when only one player is investing. By contrast, for a cooperator, their opponent will also either defect or cooperate; however, the payoff difference between these two scenarios corresponds to the second margin of the benefit function, which is diminished compared to the first. In the specialization outcome the two types experience these scenarios equally frequently, so their variance is the same when the critical point is symmetric across goods.

These results have important implications. Whereas we have seen that the exploitation outcomes produces higher fitness, we now see that the defecting type in this outcome has the highest payoff variance, which should select against defection in the presence of any demographic stochasticity (Gillespie, 1974; Constable et al., 2016; Kennedy et al., 2018; Wang et al., 2023). In a finite population, selective effects due to payoff variance will alter the expected frequency of types compared to predictions of adaptive dynamics derived in the limit of infinite population size. The effects of such stochasticity, which we predict will disrupt defectors and potentially destabilize the exploitation outcome altogether, remain an important topic for future research.

### Evolutionary instability of dimorphic critical points

For model 1, the exploitation and specialization critical points (Figure 2b, bottom left), the intermediate form of dimorphic critical point with two goods (Figure 2b, bottom right), and the dimorphic critical point in the one-good setting are all convergently stable but evolutionarily unstable. This means that investment levels in a two-type population will evolve to one of these points (i.e. convergence); at that point, one or both types can be invaded by mutants with similar investment levels. As a result, the population will undergo a second evolutionary branching, and the adaptive dynamics of investments will continue with three or four types.

These more complex divisions of labor can be inferred from the eigenvalues of the Hessian matrix of the selection gradient, evaluated at the dimorphic critical point (Leimar, 2009). Each eigenvalue is associated with one of the four investment traits. Positive eigenvalues associated with an investment trait indicate that the dimorphic type with that trait can be invaded by two new types with nearby investment levels on either side along that trait dimension. For much of the region of (*a*_1_, *a*_2_) space that implies evolutionary branching (depicted in blue in Figure 6), all eigenvalues of the selection Hessian are positive for both the exploitation and specialization dimorphisms, implying a subsequent division of labor producing four types in either case. However, for low values of *a*_1_ and *a*_2_, only the *lower* investments are associated with positive eigenvalues (depicted in orange in Figure 6). In the exploitation outcome (*a*), this means the defecting type is invasible by a pair of similar defecting types, implying three types after the second branching event. In the specialization outcome (*b*), both specialist types are invasible by types with the same upper but different lower investments, implying 4 types after the second branching event.

**Figure 6:**
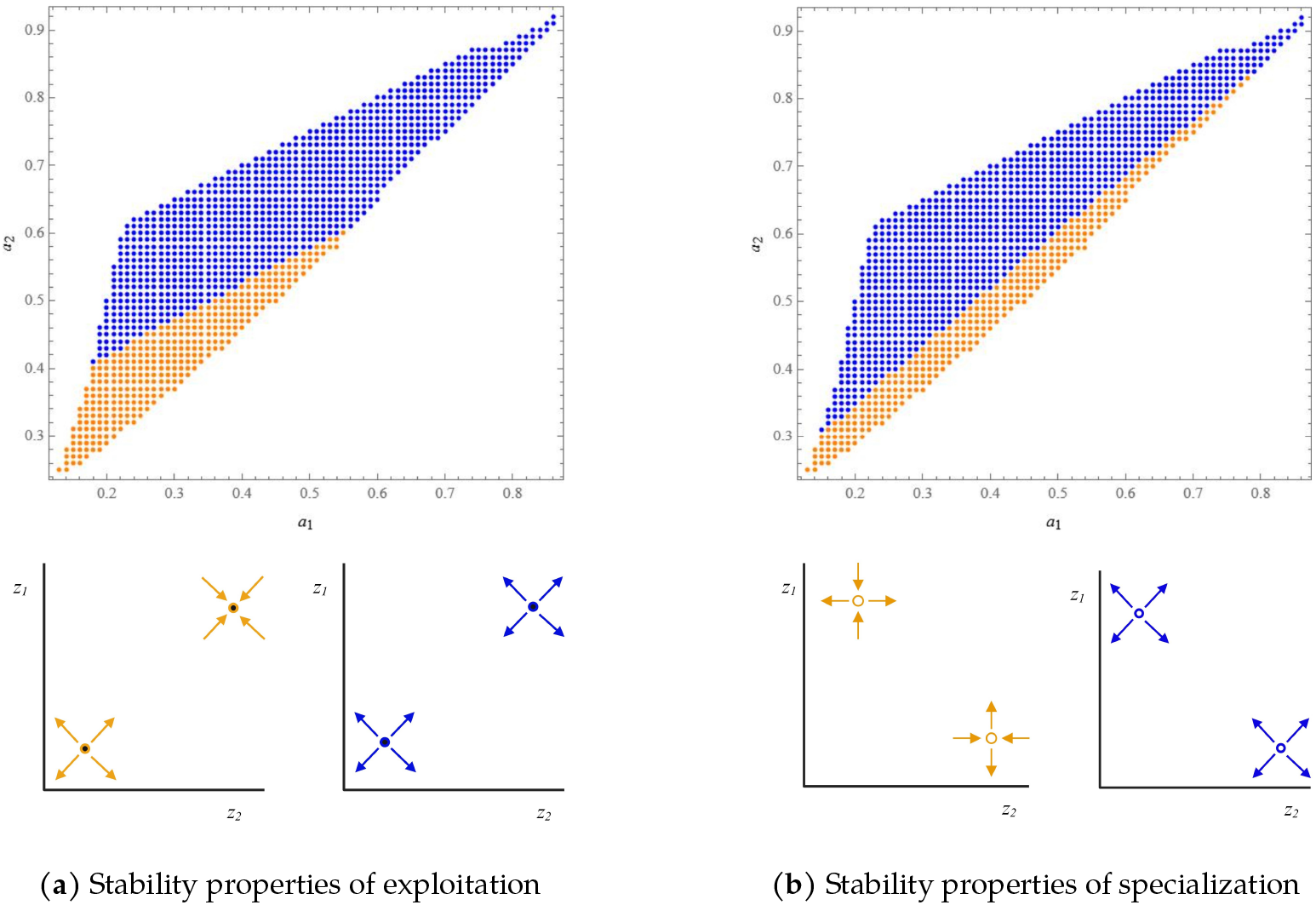
Partial or total evolutionary instability of dimorphic critical points. In the top plots, a dot with coordinates (*a*_2_, *a*_1_) indicates the evolutionary stability of the dimorphic critical point (either exploitation or specialization) with those parameters. The bottom plots illustrate expected evolutionary trajectories of investments after the dimorphic population branches, with either partial or total instability. **(Blue)** Total instability: every investment trait *z*_*u*,1_, *z*_*u*,2_, *z*_*v*,1_, *z*_*v*,2_ is unstable to invasion. **(Orange)** Partial instability: only the investment traits lower than their corresponding monomorphic critical point are unstable. In the exploitation outcome *(a)*, this means only the defecting type (with investments *z*_*v*,1_ and *z*_*v*,2_, above) is unstable. In the specialization outcome *(b)*, both types’ lower investments (*z*_*u*,2_ and *z*_*v*,1_ above) are unstable.

Similarly, in the intermediate form of division of labor for which investments in good one remain stable while branching occurs in good two, the lower investment in good two of the defecting type has a positive eigenvalue; moreover, for dimorphic populations after branching in a single public good, the lower investment of the defecting type has a positive eigenvalue. Both of these latter cases imply three types after the second branching event. Although deriving the evolutionary trajectories of the system in this more complex latter phase is beyond the scope of this paper, the important point is that subsequent branching into three or four types will assuredly occur.

Additionally, for a given parametrization of model 1, an exploitation or specialization resident population is usually invasible by types from the alternative critical point. For example, a population of specialists may be invaded by a cooperator or defector mutant; this can be thought of as a large-deviation mutation. Invasion of exploitation or specialization residents by types from the alternative division of labor was observed in a previous study (Henriques et al., 2021). This further reinforces our result that divisions of labor more complex than the dimorphic outcomes described here are the long-term outcomes of evolution in this model framework.

## Discussion

Division of labor on multiple public goods is central to the ecology of many social species (Taborsky et al., 2025). Evolutionary game theory is a natural way to model the emergence and dynamics of this phenomenon. Of particular interest is diversification of continuous investment levels in an initially-homogeneous population. Such a population finds, by incremental changes, that having two types with different effort levels brings higher expected payoff than everyone investing the same amount of effort. This emergence occurs via adaptive dynamics due to evolutionary branching of investment levels (Doebeli et al., 2004). When branching occurs in two public goods whose payoffs are added together, two basic forms of labor division, exploitation and specialization, can emerge from the same initial population (Henriques et al., 2021). In this study we have expanded our understanding of this process by generalizing the conditions for evolutionary branching in one good, analyzing the dynamics of the exploitation and specialization outcomes with two goods, and describing a novel division of labor that evolves under comparable conditions.

For many public goods ranging from extracellular enzymes in microbes to hunting for the group in humans, the public benefits and private costs vary as continuous functions of investments by individuals. The curvatures of these benefit and cost functions shape evolutionary trajectories and the resulting divisions of labor. Earlier work by Doebeli et al. (2004) described conditions for this evolutionary branching to occur under the assumption that the public benefit is a function of the sum of investments of individuals. The most important condition was that the public benefit and the private cost function both have diminishing returns (that is, a negative second derivative) with respect to investment levels. Here we have shown that this condition can be generalized to a broader family of benefit functions (11), characterized by individuals having exchangeable effects on payoffs, i.e., there are no inherent differences among individuals other than any differences in their investments. We have also shown that for exchangeable payoff functions, if 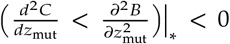, then a large enough value of *n* will ensure that the monomorphic critical point is an evolutionary branching point. Our analysis has several implications for the evolution of division of labor. First, if individuals do not differ in their competence at producing a public good, and the population initially has a monomorphic resident investment, then diminishing benefits and costs are required for the emergence of division of labor. A second requirement is that the relative rates at which benefits and costs diminish must fall within a particular range, which gets wider as *n* increases. Therefore, the best candidate populations for predicting the occurrence of division of labor will produce public goods with diminishing benefits and costs, in groups with a large (*n* > 2) number of players.

Diminishing public benefit as a function of investment has been observed across a range of species. Bacteria, for example, tend to produce extracellular metabolites at a desired concentration, above which the molecules provide less additional benefit (Foster, 2004; Gore et al., 2009; West and Cooper, 2016). Moreover, humans and other primates forage defined spaces with diminishing returns to their effort, whether or not they forage individually or collectively (Eccard and Leisenjohann, 2008). Primate hunting parties also exhibit clear diminishing returns in terms of their number, as the expected calories gained for each additional member decrease rapidly beyond some optimal size (Boesch, 1994).

Diminishing private costs may also apply for some species, if the cost per unit investment decreases due to phenotypic plasticity or a learning process. For bacteria, this is likely if the cellular machinery used to produce a given metabolite deteriorates less per molecule produced as the level of production increases. Similarly for primates and humans, private costs diminish if the physical or mental toll of a particular task decreases with the amount of effort that we put into that task.

Several studies of reproductive division of labor in single-celled organisms have found that accelerating returns to individual investments in a public good are required for labor division (Michod, 2006, 2007; West and Cooper, 2016; Cooper and West, 2018). Other studies have questioned this result, suggesting that in some cases reproductive division of labor can evolve with diminishing returns to investment, which had previously been associated with stably monomorphic contributions (Yanni et al., 2020; Cooper et al., 2021). Our present work differs in that we study division of labor in generic public goods and all individuals can reproduce as a function of their payoffs. Nonetheless, our results have parallels to those in the single-cell literature, which may help clarify the necessity of accelerating versus diminishing returns to investment. Indeed, accelerating *net* returns to a focal individual’s investment are a requirement for the emergence of division of labor via evolutionary branching (see condition (4)). The simultaneous requirement that the monomorphic critical point is convergently stable (condition (5)) implies that, when investments are exchangeable between individuals, both public benefits and private costs must diminish with respect to investments in order for division of labor to emerge (11). In cases of labor division due to accelerating returns in clonal populations, we therefore would expect to observe public benefits and private costs that diminish with respect to investments, with benefits diminishing more slowly, and the public good shared among a large number *n* of individuals. Further, if the monomorphic critical investment is evolutionarily stable (i.e. condition (4) is false) public benefits may also have diminishing returns with respect to investment, which are required if private costs are linear or diminishing (i.e. 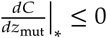).

When division of labor via evolutionary branching occurs on two public goods, several outcomes are possible. If investments in both goods would undergo branching when analyzed separately, either the exploitation or specialization division of labor may evolve, depending on initial conditions (Henriques et al., 2021). Here we have shown that members of specialization populations have lower variance in payoff but also lower fitness than members of exploitation populations. Differing payoff variance between types can reshape evolutionary dynamics due to demographic stochasticity (Wang et al., 2023). Types with low fitness can nonetheless grow in abundance, and the stability properties of critical points can be reversed compared to the no-variance setting. The fitness-variance tradeoff that we observed between the two divisions of labor could alter their dynamics in a model that incorporates stochastic effects. Moreover, it indicates a qualitative difference between the outcomes: higher fitness is achieved by cooperators and defectors via inequality of investments, while specialists achieve more reliable payoff (lower variance) by complementarity of investments. Such demographic stochasticity can select for other behavioral mechanisms such as learning to reinforce division of labor or (Netz et al., 2025). The impact of this difference on relational structure in populations is a question for future study.

In this analysis, we found that both the exploitation and specialization divisions of labor can be invaded by additional phenotypes, in two distinct ways. First, we found that the exploitation and specialization critical points are evolutionarily unstable, implying further branching events into three or four types whose investment levels evolve away from the two-type residents. Second, types from either outcome can often invade and coexist with resident populations from the alternate outcome. This matches a result of Henriques et al. (2021), who studied these outcomes using model 2 in Table 1. The implication of these results is that the long-term fate of populations in multiple goods settings is polymorphic, with a rich social structure that is unattainable in analogous models of investments in one public good.

Our adaptive dynamical analysis has several limitations. First, it applies only to continuous traits for which mutations cause a very small change in phenotype. Notably, polygenic traits fit this description, and behavioral traits such as public good production are often polygenic (Wellenreuther and Hansson, 2016). Another strong assumption made under adaptive dynamics is a haploid mode of reproduction, which can occur via either genetic inheritance or behavioral imitation (Dieckmann and Law, 1996; Metz et al., 1996). Bacteria are ideal haploid organisms for this model framework, assuming genetically inherited traits. For diploid organisms, reproductive isolation of distinct types would be required for dimorphism to persist (Palumbi, 1994). Also, this analysis applies to a behavioral trait that is learned via imitation if mutations (or “innovations”) in that trait cause very small changes. The high variability of animal behavior, especially in humans, suggests that this model framework will apply to traits learned by imitation only in exceptional circumstances.

Our results show that the long-term evolution of the division of labor can be very complex, but leave this story incomplete. We do not know if we have exhausted the list of convergently stable dimorphic critical points that are possible in this framework, or whether all such points are evolutionarily unstable. Beyond this, we do not know the evolutionary trajectories of three- and four-type divisions of labor after a secondary branching event, or whether additional polymorphic critical points exist. These questions are a fascinating topic for future research.

Our work has extended the study of division of labor on continuous investments in public goods. The results can assist researchers hoping to predict under what ecological conditions labor division is likely to occur. Beyond this, our analysis points to a vast space of complex social structures in higher-dimensional trait spaces that are evolvable by populations even under initial assumptions of total homogeneity.

## A Adaptive dynamics of investments into one public good

### A.1 Definitions

Assume there are *n* players in a public goods game, contributing investments *z*_1_, *z*_2_, …, *z*_*n*_ ∈ ℝ^+^. The public benefit *B* is a function of all investments, and the private cost *C* is a function of only the focal player’s investment. *B* and *C* are assumed to be non-negative, continuous, and increasing with the level of investment We make the consequential assumption that all players receive the same per-capita public benefit *B*, and the cost that each player pays is an identical function *C* of their own investment. The generalized payoff function from this game for player *i* ∈ (1, …, *n*) is

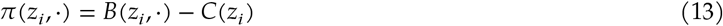

Where ⋅ denotes the *n* − 1 nonfocal investments and 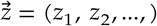.

We assume that the population is large, and that individuals play *n*-person public goods games with *n* − 1 randomly drawn others from the population. We consider a rare mutant with investment *z*_mut_, in an otherwise monomorphic population with resident investment level *z*. The mutant’s fitness is equal to the expected payoff it receives from the public goods game:

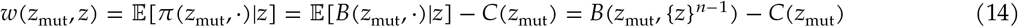

Where in the RHS, benefit is written as a function of mutant trait *z*_mut_ and {*z*}^*n*−1^ = (*z, z*, …, *z*) is a vector of length *n* − 1 with all elements equal to resident trait *z*. Below, for clarity in defining the critical point, we substitute 𝔼[*B*(*z*_mut_, ⋅)|*z*] = *B*(*z*_mut_, {*z*}^*n*−1^), which we abbreviate as *B*.

Under the framework of adaptive dynamics(Dieckmann and Law, 1996), the resident investment *z* evolves in the direction of the selection gradient:

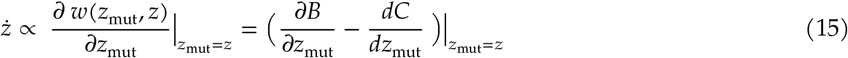

A critical point *z*_∗_ of the dynamics occurs when the selection gradient is zero:

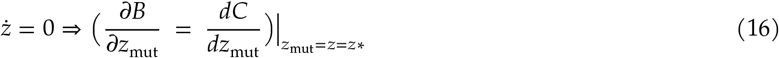

When the resident population is at the monomorphic critical point *z*_∗_, the convergence and evolutionary stability of its trait distribution are determined by the definiteness of the corresponding Hessian and Jacobian matrices of the selection gradient, respectively (Geritz et al., 1998; Leimar, 2009). Let *H* (*j*) denote the Hessian (Jacobian) matrix of the selection gradient, evaluated at *z*_∗_. For *z*_∗_ to be convergently and evolutionarily stable, it is sufficient that *H* and *j* are negative definite and necessary that they are negative semidefinite. For *z*_∗_ to be an evolutionary branching point, it is necessary that *H* is either positive (semi)definite or indefinite.

With one-dimensional investment traits, *H* has a single element

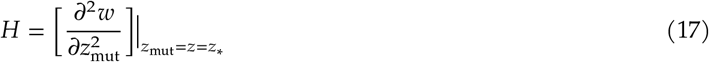

And *j* has a single element

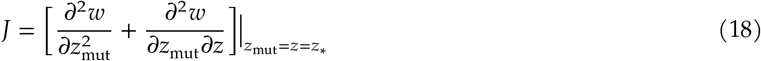

The definiteness of *H* and *j* are determined by the signs of their single elements. This gives us the following general conditions for critical point *z*_∗_ to be an evolutionary branching point:

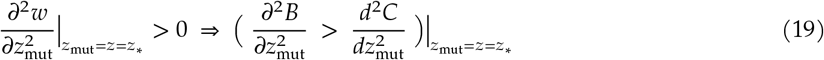

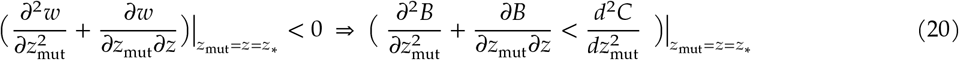

### A.2 Evolutionary branching, exchangeable payoffs, and diminishing returns

We now define the general family of ‘exchangeable’ payoff functions. Let π(*z*_*i*_, ⋅) = *B*(*f* (*z*_*i*_, ⋅)) − *C*(*z*_*i*_), where *f* (*z*_*i*_, ⋅) is an arbitrary function of all players’ investments. Label the monomorphic critical investment *z*_∗_, and compare players *i* and *j*, where *i, j* ∈ (1, …, *n*).

Assume the following are true:

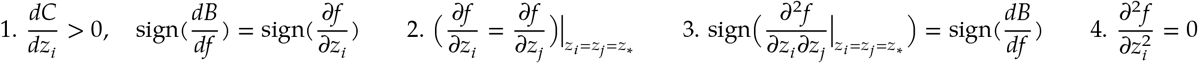

**Result 1**. *With exchangeable payoff functions as defined above, for z*_∗_ *to be an evolutionary branching point, it is necessary that* 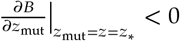 *and* 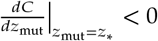

**Proof:** We first derive the the fitness of a mutant in a resident population with monomorphic trait *z*. WLOG, denote as “mut” the focal individual in the PG game, and let 2, …, *n* index the other *n* − 1 players. Then the payoff to a mutant (a random variable depending on the *n* − 1 others) is denoted π(*z*_mut_, *z*_2_, …, *z*_*n*_). The fitness of a mutant in a resident population with monomorphic trait *z* is the expected value of this RV conditioned on *z*_2_ = ⋯ = *z*_*n*_ = *z*, which is denoted *w*(*z*_mut_, *z*) = 𝔼[π(*z*_mut_, *z*_2_, …, *z*_*n*_) | *z*_2_ = ⋯ = *z*_*n*_ = *z*]. We can rewrite this as follows:

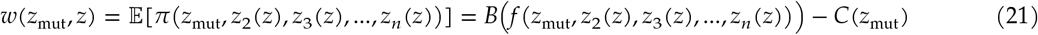

Where we define *z*_*j*_(*z*) = *z* for all *j* ∈ (2, …, *n*).

Derive (15):

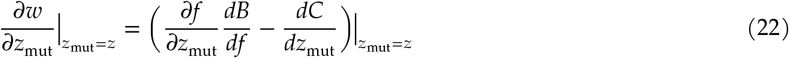

Derive (17):

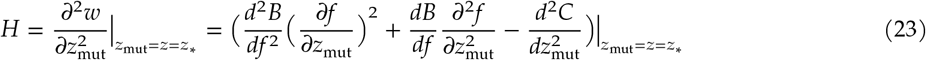

Derive (18). First, we expand the cross-derivative between the mutant and resident traits:

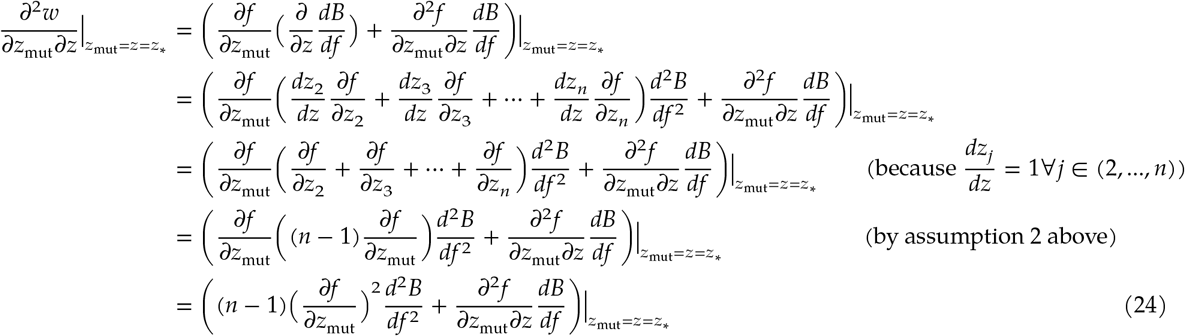

Now, we have the following expansion of (18):

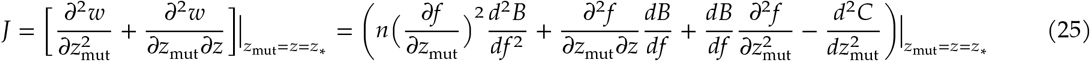

Now, simplify condition (19) above (which is (4) in main text), using derivation (23) and assumption 4 above:

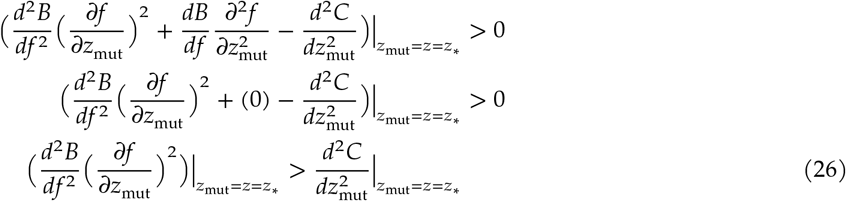

And, simplify condition (20) above (which is (5) in main text), using derivation (25) and assumption 4 above:

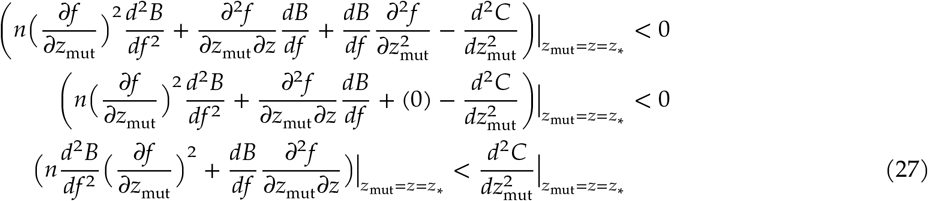

For *z*_∗_ to be an evolutionary branching point, (27) and (26) must be satisfied.

By assumption 3 above, we have that 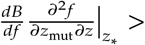 0, implying that

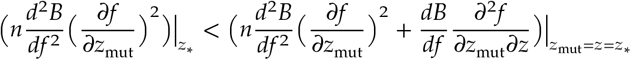

Inequality (27) therefore implies that

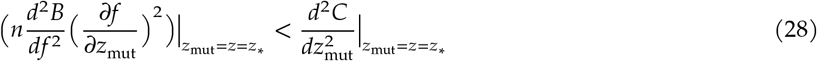

Given that *n* > 1, we infer that for (26) and (28) to both be true, it is necessary that 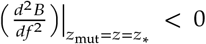.

Also, if 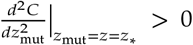, then by (26), we would have 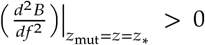, contradicting (28).

We therefore conclude that under assumptions 1–4, for *z*_∗_ to be an evolutionary branching point, it is necessary that 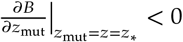 and 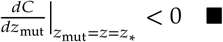

**Result 2**. With exchangeable payoff functions as defined above, 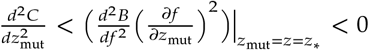, then *z* is an evolutionary branching point iff. *n* ≥ 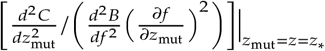.

The proof follows directly from inequalities (26) and (28). The interpretation of this result is that the volume of parameter space that implies evolutionary branching grows with increasing *n*.

Results 1 and 2 are generalizations of a result shown by (Doebeli et al., 2004) with 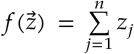 (S.I. section 1.1.3), so that public benefit is an arbitrary function of the *sum* of all *n* players’ investments (summarized by the evolutionary branching conditions in Table 1, row 2). We derive exact evolutionary branching conditions for the remaining model families in Table 1 in the following three sections.

### A.3 Evolutionary branching conditions with arbitrary multiplicative payoff functions

Here we derive evolutionary branching conditions when public benefit is an arbitrary function of the *product* of all *n* players’ investments into the public good (Table 1, row 3). In the notation above, we have 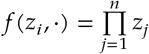.

First, write the payoff to a focal individual *i* from the public goods game with *n* − 1 other players:

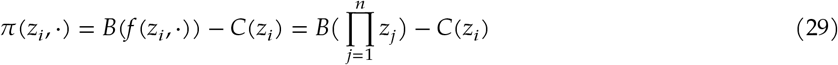

Where we make assumptions 1–4 defined in section A.2 above (note that assumptions 2 and 4 apply here by definition). We also make the underlying assumptions that *n* > 1 and that the monomorphic critical point *z*_∗_ is positive, if it exists.

Let *z* be the monomorphic resident investment level in the population, and let *z*_mut_ be a mutant trait. In this resident population we have 𝔼[*f* (*z*_mut_, ⋅)] = *z*_mut_*z*^*n*−1^ The fitness of a mutant is therefore

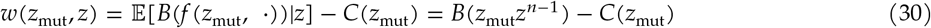

The selection gradient is

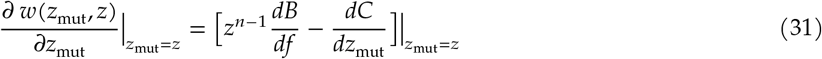

A critical point *z*_∗_ of the dynamics occurs when the selection gradient is zero:

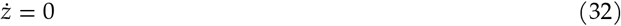

We can now use equations (17)–(20) to derive evolutionary branching conditions with multiplicative payoffs. We derive (19) as follows:

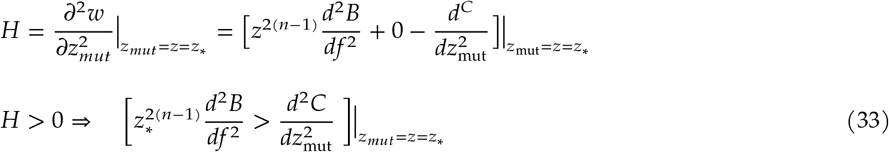

And we derive (20) as follows:

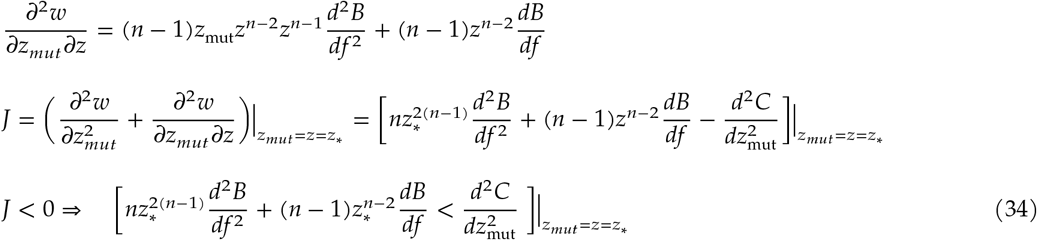

Given *n* > 1, *z*_∗_ > 0 and assumption 1 above, we infer that for (33) and (34) to both be true, it is necessary that both sides of inequality (33) be negative. Conditions (33) and (34) are reported in Table 1, row 3.

### A.4 Evolutionary branching conditions for parametric model 1

Here we derive branching conditions for model 1 (Table 1, row 4; Figure 2a, region (*ii*)) as functions of payoff parameters. We use model 1 to numerically investigate the evolution of investments into two goods.

First, write the payoff to a focal individual *i* from the public goods game with *n* − 1 other players:

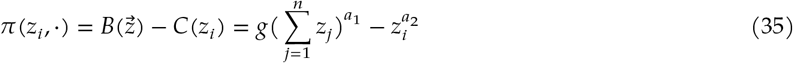

Where by assumption, *g* > 0, *a*_1_ > 0, and *a*_2_ > 0.

Next, derive the critical investment level *z*_∗_ as follows. As above, let *z* be the monomorphic resident investment level in the population, and let *z*_mut_ be a mutant trait. The fitness of a mutant is

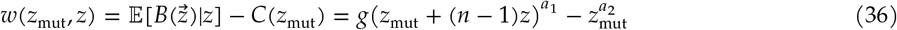

The selection gradient is

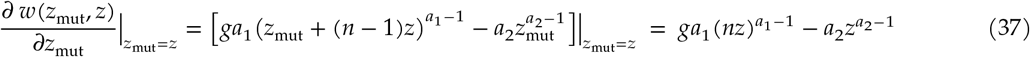

A critical point *z*_∗_ of the dynamics occurs when the selection gradient is zero:

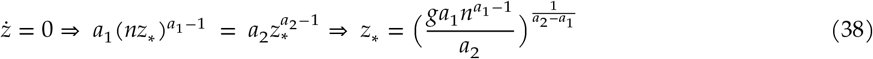

*NB: if a*_1_, *a*_2_ ∈ (0, 1), *a*_1_ < *a*_2_, *and g* > 0, *then z*_∗_ *for model 1 is strictly positive; z*_∗_ *increases with increasing a*_1_; *and z*_∗_ *decreases with increasing a*_2_. This is implied by the evolutionary branching conditions below and depicted in Figure 3 of the main text.

We can now use equations (**??**)–(20) to derive evolutionary branching conditions for model 1:

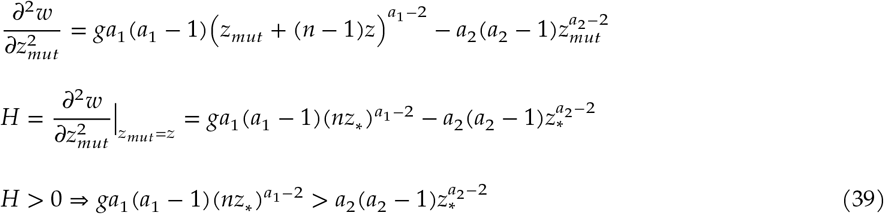

And we have

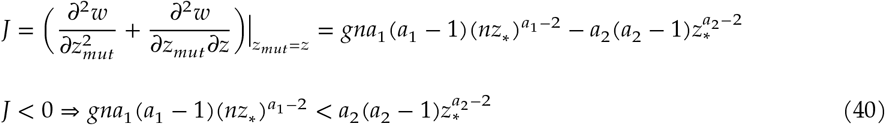

Given that *n* > 1, we infer that for (39) and (40) to both be true, it is necessary that both sides of inequality (39) be negative, that is 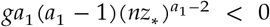 and 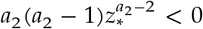. Note that we made a similar general conclusion for inequalities (26) and (28).

Because *g* > 0 and *a*_1_, *a*_2_ > 0 by assumption, (*a*_1_ − 1) and (*a*_2_ − 1) are the only terms that can be negative in (39); from this, we can conclude that *a*_1_ ∈ (0, 1) and *a*_2_ ∈ (0, 1) are necessary for evolutionary branching.

Now, we can simplify inequalities (39) and (40) as follows:

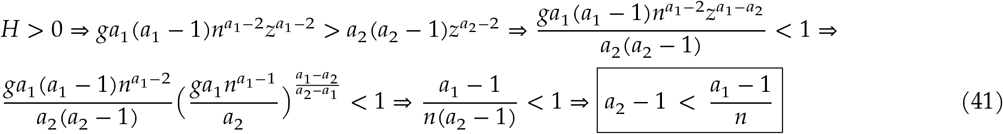

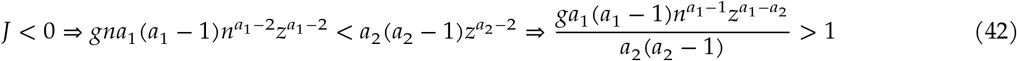

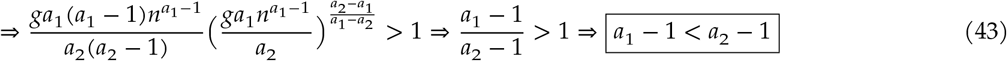

### A.5 Evolutionary branching conditions for parametric model 2

Payoff to focal *i* ∈ (1, …, *n*):

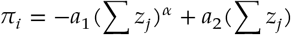

Where *α, a*_1_, and *a*_2_ are all positive.

Fitness of mutant with trait *z*_mut_ given monomorphic resident investment *z*:

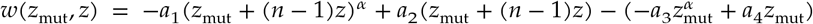

We have

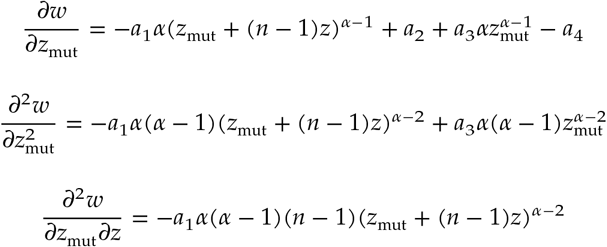

Condition (19) gives

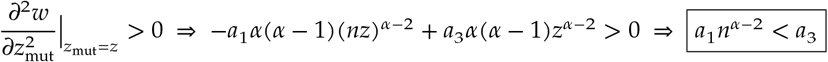

And condition (20) gives

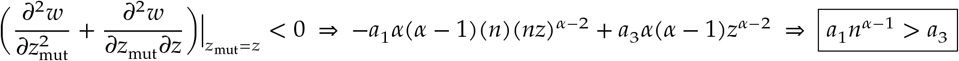

## B Adaptive dynamics of two types after evolutionary branching, with one public good

### B.1 Existence and uniqueness of frequency equilibrium *p*∗ implies unique trajectories

The deterministic expected trajectory of an evolutionary branching process (Metz et al., 1996; Geritz et al., 1998) is such that the monomorphic population evolves to a critical trait value *z*_∗_, and there changes into a dimorphic population with resident types *u* and *v* (*wlog*), with traits *z*_*u*_ > *z*_*v*_.^∗^. Let the frequency of type *u* (*v*) be *p* (1 − *p*). We denote as *w*(*z*_mut_, *z*_*u*_, *z*_*v*_, *p*) the fitness of a mutant with trait *z*_mut_ in a resident population with types *u* and *v* at relative frequency *p*. Ecological equilibrium will be reached when

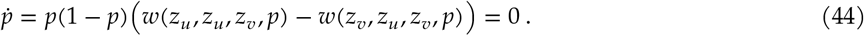

That is, the fitness of an individual of either type in the dimorphic resident population must be equal.

Solving this equation for *p* gives us a two type resident population equilibrium *p*_∗_(*z*_*u*_, *z*_*v*_) that is a function of resident traits *z*_*u*_ and *z*_*v*_. This is the derivation for (**??**) in the main text.

Adaptive dynamics proceeds by assuming that the ecological dynamics are on a shorter timescale than the evolutionary dynamics, such that in the latter timescale, the fitness of a mutant with trait *z*_mut_ in the dimorphic resident population is given by *w*(*z*_mut_, *z*_*u*_, *z*_*v*_, *p*_∗_), that is, at ecological equilibrium.

*Caveat*: it is possible that (44) has 0 or multiple solutions. The former case (0 solutions) is ruled out by the existence of evolutionary branching *only* for dimorphisms that are infinitesimal deviations from the monomorphic critical point. The latter case (multiple solutions) would imply that the ecological timescale cannot be ‘skipped over’ in deriving the dynamics, meaning an adaptive dynamical approach is not applicable.

However, result 4 below shows that for a subset of payoff functions in the Summed *z* family, including parametric models 1 and 2 (Table 1, rows 2, 4 and 5), if conditions for evolutionary branching are met and an interior frequency equilibrium *p*_∗_ ∈ (0, 1) exists, then *p*_∗_ is unique. The two-type selection gradient for resident traits is then well-defined on the subspace of (*z*_*u*_, *z*_*v*_, *p*)−space where *p* = *p*_∗_(*z*_*u*_, *z*_*v*_). This enables us to initialize a dimorphic population at a point that we ‘guess’ to be near evolutionary branching point *z*_∗_, and then derive the adaptive dynamics of *z*_*u*_ and *z*_*v*_ by integrating the two-type selection gradient.^†^

**Result 4:** *Let* 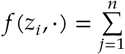 *so that the payoff function for one public good for player i* ∈ (1, …, *n*) *is* π(*z*_*i*_, ⋅) = *B*(*f* (*z*_*i*_, ⋅))− *C*(*z*_*i*_) *(with other assumptions as defined in section A*.*2), and let the investments of dimorphic types be (wlog)* 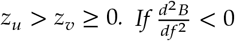 *on* ℝ^+^ *and an interior solution p*_∗_ ∈ (0, 1) *to* (44) *exists, then p*_∗_ *is unique*.

**Proof:** We follow (Peña et al., 2014) and apply their Result 3 to the gain sequence in the public goods game (defined below) to show the existence and uniqueness of *p*_∗_.

The focal individual plays with *n* − 1 others. Let *a*_*k*_ be the payoff to an individual investing *z*_*u*_ when *k* others invest *z*_*u*_ (and *n* − 1 − *k* players invest *z*_*v*_), and let *b*_*k*_ be the payoff to an individual investing *z*_*v*_ when *k* others invest *z*_*u*_. Also, define *d*_*k*_ ∶= *a*_*k*_ − *b*_*k*_ and the gain sequence **d** ∶= (*d*_0_, *d*_1_, …, *d*_*n*−1_). Also, denote the first forward difference of the gain sequence as △*d*_*k*_ = *d*_*k*+1_ − *d*_*k*_.

We have

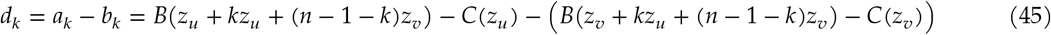

And we expand (44) as follows:

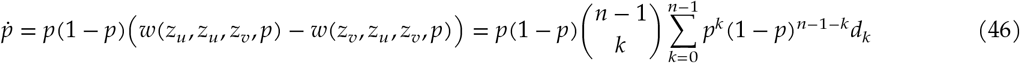

Note that (46) is equivalent to equation (1) in (Peña et al., 2014) giving the replicator dynamics of frequency *p* (which they denote *x*). Also, the fitness differential *w*(*z*_*u*_, *z*_*u*_, *z*_*v*_, *p*) −*w*(*z*_*v*_, *z*_*u*_, *z*_*v*_, *p*) is equivalent to the gain function *g*(*x*) in their equation (1), which they define in equation (2). This equivalence allows us to apply Result 3 of (Peña et al., 2014).

By Result 3 of Peña et al. (2014), if **d** has a single sign change and *d*_0_ > 0, then any interior solution *p*_∗_ ∈ (0, 1) of (44) is unique and stable under replicator dynamics (i.e. an ecological equilibrium). We prove that this is true by showing that under the assumptions above, △*d*_*k*_ is strictly negative, *d*_0_ > 0, and *d*_*n*−1_ < 0.

It turns out that if 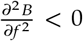 on ℝ^+^, then △*d*_*k*_ > 0, *d*_0_ < 0, and *d*_*n*−1_ < 0. This is due straightforwardly to the property of diminishing returns in *B*. We have

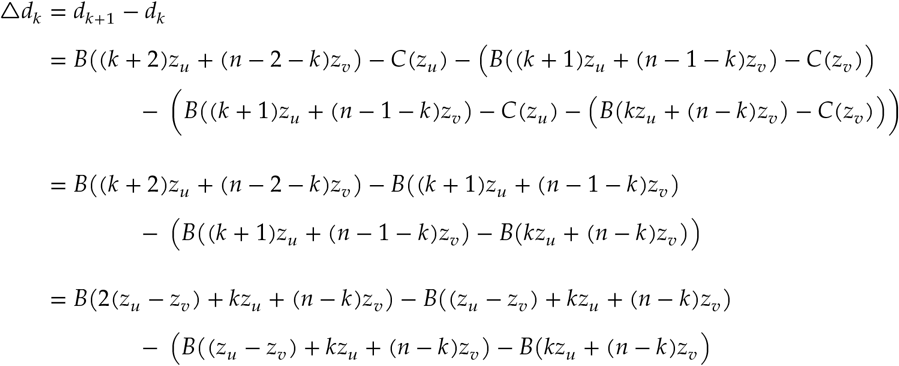

The last line above makes it clear that if 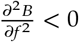 then △*d* < 0. This is because for a continuous, real-valued function *g*(*x*), for *x*_0_ < *x*_1_ < *x*_2_ with 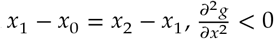 implies that *g*(*x*_2_) − *g*(*x*_1_) < *g*(*x*_1_) − *g*(*x*_0_).

We now show that if a frequency equilibrium *p*_∗_ ∈ (0, 1) exists, it is sufficient that 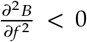 for *d*_0_ < 0 and *d*_*n*−1_ < 0 to hold. We have

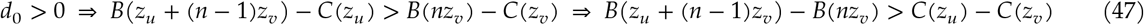

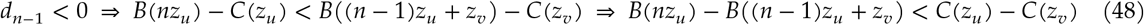

Following (44) (or (7), main text), at candidate solution *p*_∗_ we have

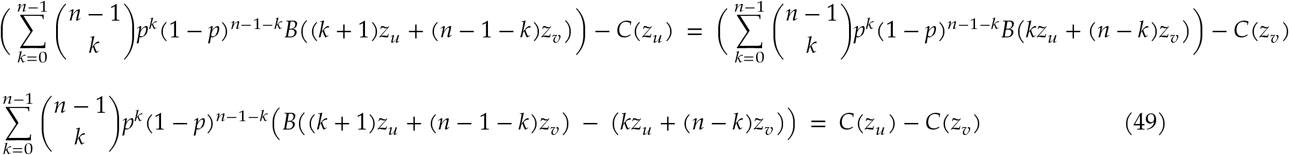

We show by contradiction that 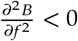 and (49) imply *d*_0_ > 0 (47) and *d*_*n*−1_ < 0 (48).

First, assume that *d*_0_ < 0, that is *B*(*z*_*u*_ + (*n* − 1)*z*_*v*_) − *B*(*nz*_*v*_) < *C*(*z* _*u*_) − *C*(*z*_*v*_). Because 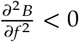, each successive term in the left-hand expected value (with *k* > 0) in (49) is smaller than the interval *B*(*z*_*u*_ + (*n* − 1)*z*_*v*_) − *B*(*nz*_*v*_). This implies that

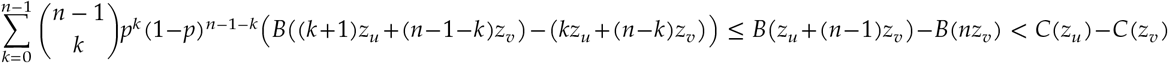

by assumption, which contradicts (49).

Similarly, assume that *d*_*n*−1_ > 0, that is *B*(*nz*_*u*_) − *B*((*n* − 1)*z*_*u*_ + *z*_*v*_) > *C*(*z*_*u*_) − *C*(*z*_*v*_). Because 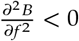, each prior term in the left-hand expected value (with *k* < *n* − 1) in (49) is larger than the interval *B*(*nz*_*u*_) − *B*((*n* − 1)*z*_*u*_ + *z*_*v*_). This implies that

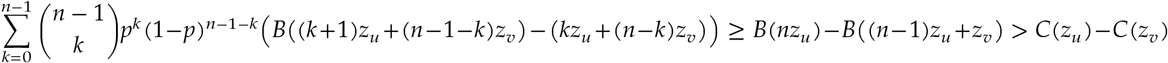

by assumption, which contradicts (49).

This completes the proof. ▪

We now are certain that for a public good whose payoff function has diminishing returns with respect to all investments on its entire domain, the occurrence of evolutionary branching leads to a dimorphic population with a unique two-dimensional adaptive dynamical trajectory.

### B.2 Definitions of two-type fitness, selection gradient, and variance in payoff

After branching, two resident types *u* and *v* co-exist in the population. Following the same derivation as for one good, fitness for a mutant with trait 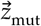 is equal to the expected payoff when its *n* − 1 opponents in the public goods game are drawn from the population with resident types *u* and *v* with probabilities *p* and 1 − *p*, respectively. That is

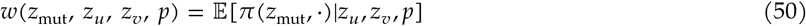

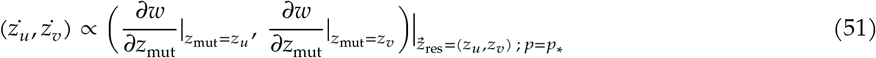

For an arbitrary individual *q* with investment *z*_*q*_ in a population with resident types *u* and *v*, we can compute *q*’s variance in payoff as follows:

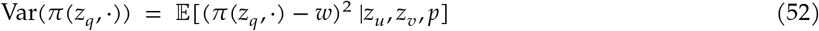

Finally, for our analyses of evolutionary (in)stability, we define the Hessian and Jacobian matrices of the selection gradient, following (Leimar, 2009):

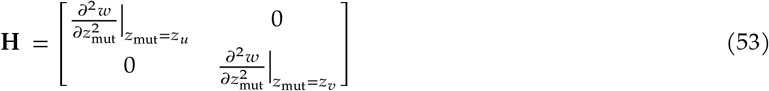

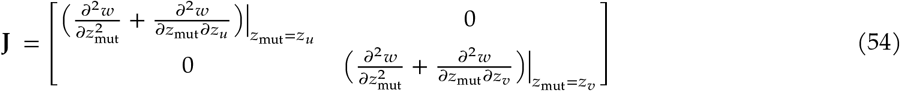

Where all elements of both matrices are evaluated at the resident dimorphic point (*z*_*u*_, *z*_*v*_, *p*_∗_).

### B.3 Numerical integration of adaptive dynamics with two types using model 1

We are now able to derive trajectories of the investments of two resident types after they emerge due to evolutionary branching. We use parametric model 1 for payoffs from the public good, which allows for numerical integration of the selection gradient. First, we choose initial dimorphic investments *z*_*u*_(*t* = 0) and *z*_*v*_(*t* = 0) that are close to the monomorphic critical investment (and evolutionary branching point) *z*_∗_. For model 1, the set of initial investments that have an interior frequency equilibrium *p*_∗_ ∈ (0, 1) is large, and includes dimorphisms that are on the same side of *z*_∗_.

Given initial investments *z*_*u*_(*t* = 0) and *z*_*v*_(*t* = 0) with *p*_∗_ ∈ (0, 1), we integrate the selection gradient (**??**) along the manifold defined by (44), on which *p* = *p*_∗_. Trajectories may either converge to a dimorphic critical point on the manifold defined by *p* = *p*_∗_, or evolve to a boundary of this manifold, at which point the two types can no longer coexist (i.e. *p*_∗_ = 0 or 1). In the former case of a dimorphic critical point, the evolutionary stability of the population must be assessed to determine whether it will remain in this configuration for infinite time or continue evolving with more types, in a three- or four-dimensional trait space.

For model 1, if (*B*_1_, *C*_1_) and (*B*_2_, *C*_2_) both meet conditions for evolutionary branching and *z*_*u*_ and *z*_*v*_ are initialized with *p* = *p*_∗_, the vast majority of trajectories will converge to a unique dimorphic critical point *z*_*v*_ < *z*_∗_ < *z*_*u*_, with the exceptions being those that are initialized near the boundary of the *p* = *p*_∗_ manifold. This allows us to readily compute the dimorphic critical point across a large region of (*a*_1_, *a*_2_) parameter space (such as in main text Figure 6 with two goods). We have made this observation numerically, because the dimorphic critical point has no closed form solution. We have not derived analytical conditions for when this point does or does not exist. Also, it is possible that an existing dimorphic critical point cannot be found, due to numerical error if the lower trait value *z*_*v*_ is very close to 0 (see the blank space on the left in main text Figure 6).

## C Evolution of division of labor on two public goods

### C.1 Definitions with two goods and two types

With two public goods, each individual now has two investment traits, such that for individual *i* ∈ (1, …, *n*), their investments are written 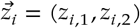.

Following (10) in main text, the simplest way to define the payoff from two public goods is as follows. Let *B*_*k*_ and *C*_*k*_ denote the benefit and cost functions for good *k*, and let {*z*_−*i,k*_} denote the *n* − 1 nonfocal investments in good *k*. Payoff for *i* is

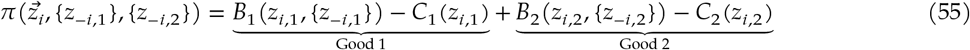

We label the two types that arise after branching *u* and *v*, with investment traits 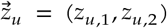 and 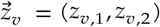 and frequencies *p*, (1 − *p*). As above, the fitness of a mutant with trait vector 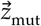 in this population is

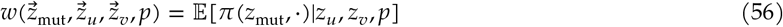

The selection gradient 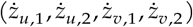 for each of the four investment traits is proportional to

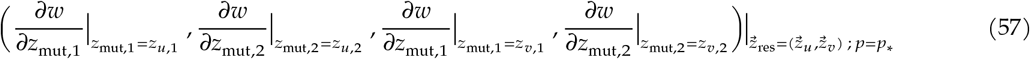

Where *p* = *p*_∗_ is the solution of 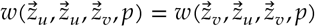, as in (44).

The variance in payoff experienced by individuals with arbitrary trait vector 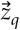 in the dimorphic resident population is defined as

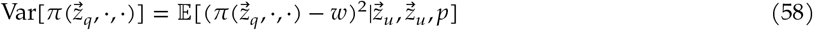

Finally, following Leimar (2009) as before (see section **??**), we define the Hessian and Jacobian matrices of the selection gradient for two goods and two types.

The Hessian matrix is block diagonal with entries equal to the second partial derivatives of fitness with respect to both mutant traits, evaluated at the resident point. Following equation (7) from (Leimar, 2009), we can denote one of these blocks for type *k* ∈ {*u, v*} as follows:

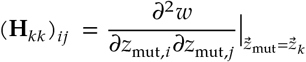

Where the first (second) partial derivative is taken with respect to a mutant of type *k*’s investment in public good *i* (*j*), with *i, j* ∈ {1, 2}. Note that because in our payoff function investments in different goods affect different summands (i.e. there are no interactions between goods), we know that (**H**_*kk*_)_*ij*_ = 0 for *i* ≠ *j*.

This gives us the following Hessian matrix for two goods and two types (using column ordering (*z*_*u*,1_, *z*_*u*,2_, *z*_*v*,1_, *z*_*v*,2_)):

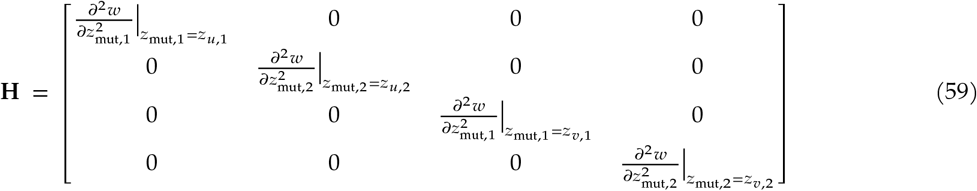

Where, as before with one good, all elements of *H* are evaluated at the resident dimorphic point 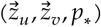.

To define the Jacobian, we refer to equation (10) in (Leimar, 2009), which defines the matrix **Q** of cross-derivatives of fitness with respect to all mutant traits and all resident traits. We can denote a block of such cross derivatives where the mutant is from type *k* and the resident is from type *l* as follows:

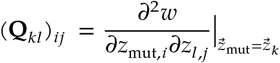

The desired Jacobian matrix is evaluated **J** = **H+Q**. We write this 4 × 4 matrix below in two blocks: the left half (4 × 2) and the right half (4 × 2).

**J** = **H+Q** =

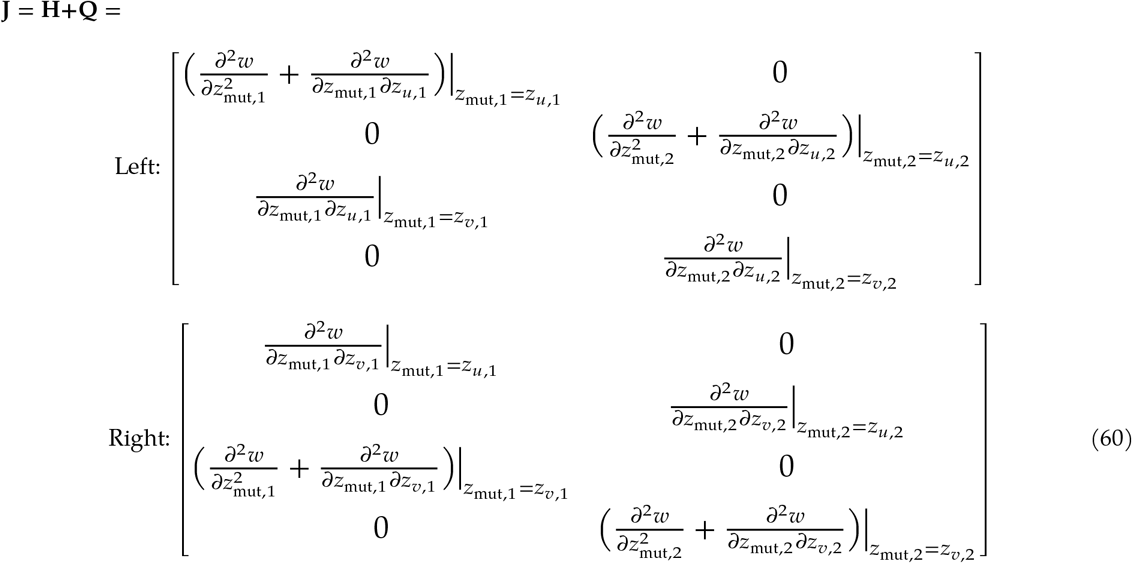

### C.2 Numerical integration of dynamics with two goods and two types, using model 1

We numerically derive adaptive dynamical trajectories of investments with two public goods and two types analogously to our approach with one good and two types (see section B.3), by initializing two types with investment traits 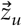 and 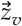 near the evolutionary branching point 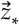 and integrating (57). This approach allows us to confirm the characteristics of dimorphic critical points shown in Figure 2b (main text), and their relationship to the stability properties of their prior monomorphic critical points, indicated by Figure 2a.

Results in main text Figures 3–6 are reported with *n* = 2 and *B*_1_ = *B*_2_, *C*_1_ = *C*_2_ at the respective dimorphic critical points for the exploitation and specialization outcomes. For model 1, the symmetry across goods implies *g*_1_ = *g*_2_, *a*_1_ = *a*_3_, and *a*_2_ = *a*_4_. We found results to be qualitatively the same for *n* = 3 (within the appropriate region of parameter space). Also, results with small asymmetry across goods (that is, small values of |*g*_1_ − *g*_2_|, |*a*_1_ − *a*_3_|, and |*a*_2_ − *a*_4_|) are qualitatively similar, though asymmetry leads to a larger set of discontinuities on the manifold defined by *p* = *p*_∗_ (Eq. (44)).

### C.3 Adaptive dynamics with two public goods and two types after evolutionary branching

If (4) and (5) are true for either good 1 (*B*_1_, *C*_1_) or good 2 (*B*_2_, *C*_2_), then evolutionary branching into two types will occur. For payoff model 1, if (19) and (20) are true for both goods, we have found that the two types evolve to one of two (or four) dimorphic critical points. These points represent two characteristically different divisions of labor: exploitation (ex), in which some individuals invest more in both goods while others invest less in both; and specialization (sp), in which some individuals invest more in good 1 and less in good 2, while others invest more in good 2 and less in good 1 (Figure 2b). The exploitation and specialization outcomes are multistable. In our numerics, the following basins for these outcomes hold consistently:

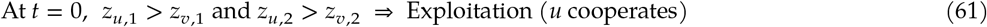

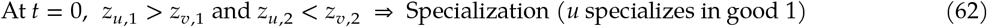

However, we have not analytically proved that these basin definitions are always true in this setting.

If (4) and (5) are true for one good while the monomorphic critical point for the other good is evolutionarily stable (see main text Figure 2b, bottom right), the population branches into two types which invest divergent amounts in the former but equal amounts in the latter (stable) good. Trajectories in this intermediate regime evolve to a dimorphic critical point less consistently than the exploitation–specialization regime above, because the manifold defined by *p* = *p*_∗_ (44) has a larger set of discontinuities. We are able to compute some consistent qualitative characteristics of this division of labor, such as variance in payoff for types and evolutionary stability (described below).

### C.4 Bifurcation in investment levels for exploitation and specialization

The characteristics of the dimorphic critical points corresponding to exploitation and specialization divisions of labor are shown in Figure 3 (main text). As described, when payoff parameters are symmetric across the two goods, there is a bifurcation as *a*_2_ increases above *a*_1_. Before the threshold value of *a*_2_, there are two dimorphic critical points with symmetric investments across goods: for exploitation *z*_*v*1_ = *z*_*v*,2_ < *z*_∗_ < *z*_*u*,1_ = *z*_*u*,2_, whereas for specialization *z*_*v*_ = *z*_*v*,2_ < *z*_∗_ < *z*_*u*,1_ = *z*_*u*,2_, where *z*_∗_ is the common monomorphic critical point for both goods. Note that across these outcomes, the upper and lower investments need not be equal. In the specialization case, symmetry of investments across goods constrains the frequencies of the types to be equal, that is *p*_∗_ = 0.5.

When *a*_2_ sufficiently exceeds *a*_1_, the specialization outcome has four additional dimorphic critical points that have asymmetric investments across goods 1 and 2: (*z*_*v*,1_ < *z*_*u*,2_) < *z*_∗_ < (*z*_*u*,1_ < *z*_*v*,2_) (Figure 3b, right side) ; (*z*_*u*,2_ < *z*_*v*,1_) < *z*_∗_ < (*z*_*v*,2_ < *z*_*u*,1_); (*z*_*v*,2_ < *z*_*u*,1_) < *z*_∗_ < (*z*_*u*,2_ < *z*_*v*,1_); and (*z*_*u*,1_ < *z*_*v*,2_) < *z*_∗_ < (*z*_*v*,1_ < *z*_*u*,2_). These four critical points are multistable with the symmetric points *z*_*v*1_ = *z*_*v*,2_ < *z*_∗_ < *z*_*u*,1_ = *z*_*u*,2_ and *z*_*u*1_ = *z*_*u*,2_ < *z*_∗_ < *z*_*v*,1_ = *z*_*v*,2_. However, we have observed that past the bifurcation point, the basins of attraction for the latter symmetric critical points are marginal neighborhoods around those points, while the corresponding basins for the asymmetric critical points are much larger (that is, most of the *p* = *p*_∗_ manifold).

When payoff parameters are asymmetric across the two goods, qualitative results are similar to the specialization outcomes with asymmetric investments that occur when payoff parameters are symmetric across goods. Results concerning fitness of outcomes, variance in payoff across types and outcomes, and evolutionary stability hold when payoff parameters are asymmetric across the two goods, for small values of |*g*_1_ −*g*_2_|, |*a*_1_ −*a*_3_|, and |*a*_2_ −*a*_4_|. However, when payoff parameters for the two public goods are sufficiently different, these relationships may change. Because of this and the increased complexity of the *p* = *p*_∗_ manifold in the asymmetric-goods setting, we report comparisons between exploitation and specialization outcomes in the symmetric-goods setting.

### C.5 Computation of fitness and variance in payoff

Under adaptive dynamics, the fitnesses of both types at any dimorphic point are equal, that is 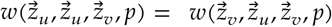 44. We computed this fitness level for the exploitation (*w*_ex_) and specialization (*w*_sp_) outcomes and compared these two levels in Figure 4 (main text) across the appropriate parameter space. We consistently found that the maximum value of *w*_ex_ − *w*_sp_ occurred at the lowest value of *a*_2_ that exceeded the bifurcation threshold in the specialization outcome.

We compute the payoff variance for types and the mean payoff variance for outcomes using (52) (or (9) in main text) with one public good and (58) with two public goods. The overall result is that in a dimorphic population after evolutionary branching, the type with lower investment level(s) has higher payoff variance. We have not proven that this always holds for dimorphic critical points. As mentioned in the main text, differential variance may be consequential for dynamics in the presence of demographic stochasticity.

### C.6 Evolutionary instability of dimorphic critical points

We analyze the Hessian (see (53) and (59)) and the Jacobian (see (54) and (60)) of the selection gradient for dimorphic critical points for payoff model 1 with one and two public goods, including in the latter case the exploitation, specialization, and intermediate divisions of labor.

The selection Jacobian is negative definite for every critical point we have observed, indicating that each of these points is convergently stable. We have verified numerically by repeated integration that the neighborhood of convergence stability for each dimorphic critical point includes initial dimorphisms that are small deviations from the evolutionary branching point. This further confirms that population will evolve to the dimorphic critical point after branching.

However, as reported in the main text, all dimorphic critical points are evolutionarily unstable due to either a positive definite or indefinite selection Hessian. Each trait (*z*_*u*,1_, *z*_*u*,2_, *z*_*v*,1_, *z*_*v*,2_) in the population has a straightforward mapping to one of the eigenvalues of **H**, allowing us to infer which of the traits are evolutionarily unstable in the case when **H** is indefinite. The eigenvectors of **H** form the 4 × 4 identity matrix, that is *v*_1_ = (1, 0, 0, 0)^*T*^, *v*_2_ = (0, 1, 0, 0)^*T*^, and so on. We can therefore map each eigenvector to one of (*z*_*u*,1_, *z*_*u*,2_, *z*_*v*,1_, *z*_*v*,2_), in that order (defined by **H**). Of course, there is an eigenvalue associated with each eigenvector, and thus with each of the four trait dimensions. If **H** is indefinite, some eigenvalues are positive while the others are negative. We infer that the investment traits that are associated with positive eigenvalues are evolutionarily unstable, and can be invaded by mutants with slightly different investments along that trait dimension; conversely, investment traits that are associated with negative eigenvalues are evolutionarily stable, and cannot be invaded by nearby mutants. The results concerning evolutionary instability and corresponding inferences about higher-dimensional populations are summarized in “Evolutionary instability of dimorphic critical points”, main text.

### C.7 Mutual invasibility of exploitation and specialization divisions of labor

We found that the exploitation division of labor was often invasible by types from the specialization outcome for the same parametrization, and vice versa. We checked this by computing the mutant fitness of types in their alternative divisions of labor. For clarity, denote the type *u* and *v* trait vectors and frequency equilibrium *p*_∗_ with a second subscript, ‘ex’ or ‘sp’, indicating the division of labor outcome. We checked these four inequalities:

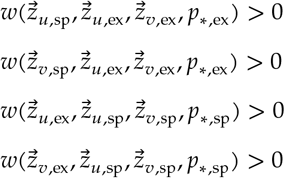

If any of these inequalities are true, the associated mutant type can invade the alternative division of labor; for example if the first inequality is true, the *u* type from the specialization outcome can invade the exploitation division of labor with the same parameters.

In general, some or all of these inequalities are true for each parametrization in the region that determines evolutionary branching. Analytical conditions for when each inequality is true are beyond the scope of this paper.

### C.8 Comparison to results with payoff model 2

Model 2 (Table 1, row 5) above was investigated by (Henriques et al., 2021), who observed the same multistability between the exploitation and specialization divisions of labor. For model 2, the *B* and *C* functions are quadratic; beyond their maxima, they decrease with investments. As a result, exploitation and specialization outcomes with model 2 correspond to constrained endpoints 0 and min(max(*B*),max(*C*)) of the dynamics, rather than critical points.

We computed fitness, population-level variance in payoff, and type-level variance in payoff at the exploitation and specialization endpoints with model 2, across a range of parameters that determine evolutionary branching. Comparing these results with those from model 1 offers some additional insight. First, fitness in an exploitation population can be lower than fitness in a specialization population for model 2, unlike for model 1. Second, payoff variance for the cooperator type in the exploitation outcome is consistently lower than payoff variance for specialized types in the specialization outcome for model 2, while both are lower than the variance of defectors. For model 1, the variance for cooperators eventually exceeds that of specialists as the concavity of the benefit function increases (see main text Figure 5, row 1). Finally, one notable relationship is the same for models 1 and 2: population-level variance is higher for individuals in an exploitation population than for those in a specialization population.

## Footnotes

† Note that we do not include all terms in the canonical equation of adaptive dynamics here; because we assume no covariance between mutations in any separate traits (Leimar, 2009), we neglect these terms in describing the shape of trajectories.

∗ If *z*_*u*_ and *z*_*v*_ are the dimorphic residents *immediately* after branching, there are two possibilities: *z*_*u*_ = *z*_∗_ > *z*_*v*_ or *z*_*u*_ > *z*_*v*_ = *z*_∗_. Because the selection gradient is 0 at *z*_∗_, it contains no information about which of these outcomes is more likely to occur, as noted by (**?**).

## References

Akçay, E. and Van Cleve, J. (2012). Behavioral responses in structured populations pave the way to group optimality. American Naturalist, 179(2):257–269.

Anderson, A., Chilczuk, S., Nelson, K., Ruther, R., and Wall-Scheffler, C. (2023). The myth of man the hunter: women’s contribution to the hunt across ethnographic contexts. PLOS One, 18(6).

Apicella, C. and Silk, J. (2019). The evolution of human cooperation. Current Biology, 29:447–450.

Archetti, M. and Scheuring, I. (2012). Review: Game theory of public goods in one-shot social dilemmas without assortment. Journal of Theoretical Biology, 299:9–20.

Axelrod, R. and Hamilton, W. (1981). The evolution of cooperation. Science, 211(4489):1390–1396.

Bird, R. (1999). Cooperation and conflict: The behavioral ecology of the sexual division of labor. Evolutionary Anthropology, 8(2):65–75.

Boesch, C. (1994). Cooperative hunting in wild chimpanzees. Animal Behaviour, 48:653–657.

Constable, G., Rogers, T., McKane, A., and Tarnita, C. (2016). Demographic noise can reverse the direction of deterministic selection. PNAS, 113:4745–4754.

Cooper, G., Frost, H., Liu, M., and West, S. (2021). Does the evolution of division of labour require accelerating returns to individual specialization? biorxiv.

Cooper, G. A. and West, S. A. (2018). Division of labour and the evolution of extreme specialization. Nature Ecology and Evolution, 2:1161–1167.

Cordero, O., Ventouras, L.-A., Delong, E., and Polz, M. (2012). Public good dynamics drive evolution of iron acquisition strategies in natural bacterioplankton populations. PNAS, 109(49):20059–20064.

Dieckmann, U. and Law, R. (1996). The dynamical theory of coevolution: A derivation from stochastic ecological processes. Journal of Mathematical Biology, 34:579–612.

Diekmann, A. (1985). Volunteer’s dilemma. Journal of conflict resolution, 29(4):605–610.

Doebeli, M., Hauert, C., and Killingback, T. (2004). The evolutionary origin of cooperators and defectors. Science, 306(5964):859–862.

Drescher, K., Nadell, C. D., Stone, H. A., Wingreen, N. S., and Bassier, B. L. (2014). Solutions to the public goods dilemma in bacterial biofilms. Current Biology, 24:50–55.

Eccard, J. and Leisenjohann, T. (2008). Foraging decisions in risk-uniform landscapes. PLOS One, 3(10).

Foster, K. (2004). Diminishing returns in social evolution: the not-so-tragic commons. Journal of Evolutionary Biology, 17:1058–1072.

Frank, S. (1995). Mutual policing and repression of competition in the evolution of cooperative groups. Nature, 377:520–522.

Frank, S. (2010). A general model of the public goods dilemma. Journal of Evolutionary Biology, 23(6):1245–1250.

Geritz, S. A., Kisdi, E., MeszéNA, G., and Metz, J. A. (1998). Evolutionarily singular strategies and the adaptive growth and branching of the evolutionary tree. Evolutionary ecology, 12:35–57.

Gillespie, J. (1974). Natural selection for within generation variance in offspring number. Genetics, 76:601–606.

Gillespie, J. (1975). Natural selection for within generation variance in offspring number ii. discrete haploid models. Genetics, 81:403–413.

Gore, J., Youk, H., and Van Oudenaarden, A. (2009). Snowdrift game dynamics and facultative cheating in yeast. Nature, 459(7244):253–256.

Hamilton, W. (1964). The genetical evolution of social behavior. Journal of Theoretical Biology, 7(1):1–52.

Hauert, C. and Doebeli, M. (2004). Spatial structure often inhibits the evolution of cooperation in the snowdrift game. Nature, 428(6983):643–646.

Hauert, C., Holmes, M., and Doebeli, M. (2007). Evolutionary games and population dynamics: Maintenance of cooperation in public goods games. Proceedings of the Royal Academy B: Biological Sciences, 273(1600):2565–2571.

Henriques, G., Ito, K., Hauert, C., and Doebeli, M. (2021). On the importance of evolving phenotype distributions on evolutionary diversification. PLOS Computational Biology, 17(2).

Hooper, P., Demps, K., Gurven, M., Gerkey, D., and Kaplan, H. (2015). Skills, division of labour and economies of scale among amazonian hunters and south indian honey collectors. Phil. Trans. R. Soc., 370.

Kaznatcheev, A., Vandervelde, R., Scott, J., and Basanta, D. (2017). Cancer treatment scheduling and dynamic heterogeneity in social dilemmas of tumour acidity and vasculature. British Journal of Cancer, 116(6):785–792.

Kennedy, P., Higginson, A. D., Radford, A. N., and Sumner, S. (2018). Altruism in a volatile world. Nature, 555(7696):359–362.

Kramer, J., Özkaya, Ö., and Kummerli, R. (2019). Bacterial siderophores in community and host interactions. Nature Reviews Microbiology, 18(3):152–163.

Leimar, O. (2009). Multidimensional convergence stability. Evolutiuonary Ecology Research, 11:191–208.

Maynard Smith, J. and Price, G. (1973). The logic of animal conflict. Nature, 246:15–18.

Metz, J., Geritz, S., Meszena, G., Jacobs, S., and Heerwaarden, J. V. (1996). Adaptive dynamics: a geometrical study of the consequences of nearly-faithful reproduction. Stochastic and Spatial Structures of Dynamical Systems, 45:183–231.

Michod, R. (2006). The group covariance effect and fitness trade-offs during evolutionary transitions in individu-ality. PNAS, 103(24):9113–9117.

Michod, R. (2007). Evolution of individuality during the transition from unicellular to multicellular life. PNAS, 104(1):8613–8618.

Motro, U. (1991). Cooperation and defection: Playing the field and the ess. Journal of Theoretical Biology, 151(2):145– 154.

Netz, C., Fawcett, T. W., Higginson, A. D., Taborsky, M., and Taborsky, B. (2025). Group size and labour demands determine division of labour as a consequence of demographic stochasticity. Philosophical Transactions B, 380(1922):20240206.

Nowak, M. (2006). Five rules for the evolution of cooperation. Science, 314:1560–1563.

Oliviera, N. M., Niehus, R., and Foster, K. R. (2014). Evolutionary limits to cooperation in microbial communities. PNAS, 111(50):17941–17946.

Özkaya, Ö., Xavier, K., Dionisio, F., and Balbontin, R. (2017). Maintenance of microbial cooperation mediated by public goods in single- and multiple-trait scenarios. Journal of Bacteriology, 199(22).

Palumbi, S. (1994). Genetic divergence, reproductive isolation, and marine speciation. Annual Review of Ecological Systems, 25:547–572.

Peña, J., Lehmann, L., and Nöldeke, G. (2014). Gains from switching and evolutionary stability in multi-player matrix games. Journal of Theoretical Biology, 346:23–33.

Rankin, D., Bargum, K., and Kokko, H. (2007). The tragedy of the commons in evolutionary biology. Trends in Ecology and Evolution, 22(12):643–651.

Ross-Gillespie, A., Dumas, Z., and Kummerli, R. (2014). Evolutionary dynamics of interlinked public goods traits: an experimental study of siderophore production in pseudomonas aeruginosa. Journal of Evolutionary Biology, 28:29–39.

Rossetti, V., Schirrmeister, B., Bernasconi, M., and Bagheri, H. (2010). The evolutionary path to terminal differentiation and division of labor in cyanobacteria. Journal of Theoretical Biology, 262(1):23–34.

Smith, P. and Schuster, M. (2019). Public goods and cheating in microbes. Current Biology, 29:442–447.

Souza, M. O., Pacheco, J. M., and Santos, F. C. (2009). Evolution of cooperation under n-person snowdrift games. Journal of Theoretical Biology, 260:581–588.

Stibbard-Hawkes, D., Smith, K., and Apicella, C. (2022). Why hunt? why gather? why share? hadza assessments of foraging and food sharing motive. Evolution and Human Behavior, 43(3):257–272.

Taborsky, M., Fewell, J. H., Gilles, R., and Taborsky, B. (2025). Division of labour as key driver of social evolution. Philosophical Transactions B, 380(1922):20230261.

Wang, G., Su, Q., Wang, L., and Plotkin, J. (2023). Reproductive variance can drive behavioral dynamics. PNAS, 120(12).

Wellenreuther, M. and Hansson, B. (2016). Detecting polygenic evolution: problems, pitfalls and promises. Trends in Genetics, 32(3):155–164.

West, S. A. and Cooper, G. A. (2016). Division of labour in microorganisms: an evolutionary perspective. Nature Reviews Micriobiology, 14(11):716–723.

Yanni, D., Jacobeen, S., Marquez-Zacarias, P., Weitz, J., Ratcliff, W., and Yunker, P. (2020). Topological constraints in early multicellularity favor reproductive division of labor. eLife, 9:e54348.

Zheng, D. F., Yin, H. P., Chan, C. H., and Hui, P. M. (2007). Cooperative behavior in a model of evolutionary snowdrift games with n-person interactions. Europhysics Letters, 80(1).

